# Analytical treatment interruption after short-term anti-retroviral therapy in a postnatally SHIV infected infant rhesus macaque model

**DOI:** 10.1101/667139

**Authors:** Ria Goswami, Ashley N. Nelson, Joshua J. Tu, Maria Dennis, Liqi Feng, Amit Kumar, Jesse Mangold, Riley J. Mangan, Cameron Mattingly, Alan D. Curtis, Veronica Obregon-Perko, Maud Mavigner, Justin Pollara, George M. Shaw, Katharine J. Bar, Ann Chahroudi, Kristina De Paris, Cliburn Chan, Koen K.A. Van Rompay, Sallie R. Permar

**Author notes:** Address correspondence to Sallie R. Permar. R.G. and A.N.N. contributed equally to this work.

## Abstract

To achieve long-term viral remission in HIV-infected children, novel strategies beyond early anti-retroviral therapy (ART) will be necessary. Identifying clinical predictors of time to viral rebound upon ART interruption will streamline the development of novel therapeutic strategies and accelerate their evaluation in clinical trials. However, identification of these biomarkers is logistically challenging in infants, due to sampling limitations and potential risks of treatment interruption. To facilitate identification of biomarkers predicting viral rebound, we have developed an infant rhesus macaque (RM) model of oral SHIV.CH505.375H.dCT challenge and analytical treatment interruption (ATI) after short-term ART. We used this model to characterize SHIV replication kinetics and virus-specific immune responses during short-term ART or post-ATI and demonstrated plasma viral rebound in 5 out of 6 (83%) infants. We observed a decline in humoral immune responses and partial dampening of systemic immune activation upon initiation of ART in these infants. Furthermore, we documented that infant and adult macaques have similar SHIV replication and rebound kinetics and equally potent virus-specific humoral immune responses. Finally, we validated our models by confirming a well-established correlate of time to viral rebound, namely pre-ART plasma viral load, as well as identified additional potential humoral immune correlates. Thus, this model of infant ART and viral rebound can be used and further optimized to define biomarkers of viral rebound following long-term ART as well as to pre-clinically assess novel therapies to achieve a pediatric HIV functional cure.

**IMPORTANCE:** Novel interventions that do not rely on daily adherence to ART are needed to achieve sustained viral remission for perinatally infected children who currently rely on lifelong ART. Considering the risks and expense associated with ART-interruption trials, identification of biomarkers of viral rebound will prioritize promising therapeutic intervention strategies, including anti-HIV Env protein therapeutics. However, comprehensive studies to identify those biomarkers are logistically challenging in human infants, demanding the need for relevant non-human primate models of HIV rebound. In this study, we developed an infant RM model of oral Simian/Human Immunodeficiency virus infection expressing clade C HIV Env, and short-term ART followed by ATI, longitudinally characterizing immune responses to viral infection during ART and post-ATI. Additionally, we compared this infant RM model to an analogous adult RM rebound model and identified virologic and immunologic correlates of time to viral rebound post-ATI.

## INTRODUCTION

Despite the widespread availability and effectiveness of antiretroviral therapy (ART), each year >180,000 infants continue to become infected with HIV (1). Acquiring HIV at this early age commits these children to life-long ART, since stopping therapy is universally associated with viral rebound. However, continuous access to ART can be challenging in resource-limited settings (2), leading to treatment interruption and poor clinical outcomes. Maintaining adherence to life-long therapy is particularly challenging among adolescents (3), resulting in the development of drug resistant viral strains (4). Even if adherence is maintained, chronic exposure to ART from a young age predisposes children to drug-associated metabolic complications (5). Therefore, novel intervention strategies that do not rely on daily ART will be needed for sustained viral remission in infected children. While the establishment of viral reservoirs may not be prevented even when ART is initiated within hours of HIV infection (6), a reduced size of the latent reservoir has been demonstrated to lengthen time to viral rebound in clinical trials (7–9). Therefore, reducing the size of the viral reservoir and attaining sustained viral remission after treatment discontinuation has been the focus of an emerging global effort aimed at developing a cure for HIV infection.

As new therapeutic interventions to attain drug-free viral remission are developed and assessed in clinical trials, safe means to measure their efficacy will be needed. While mathematical models to predict viral rebound time from reservoir size have been developed (10–12), this approach is limited by the inaccuracy of existing assays to measure viral reservoir size (13) and inter-patient variability in response to identical treatment strategies. Therefore, careful monitoring of viral rebound after analytical treatment interruption (ATI) still remains the “gold standard” to accurately validate the efficacy of any novel HIV therapeutic strategy. However, this approach is logistically challenging and carries considerable risk of virus transmission and replenishment of viral reservoir upon reactivation. More importantly, this strategy will be ethically challenging in HIV-infected children, since the outcome of ATI studies on long-term pediatric health are not known. Considering these risks, identification of biomarkers to serve as predictors of time to HIV rebound (14) would be useful to prioritize development of treatment strategies, avoiding the cost and risk of ATI studies that are unlikely to have clinical efficacy.

Virologic and immunologic biomarkers predicting HIV rebound have been identified by several studies in recent years (15–18). Yet, our understanding of the predictors of HIV rebound in the setting of maturing infant immune systems is limited. These types of comprehensive studies are further complicated in infants due to limited volumes of samples that can be collected at this age. Thus, pediatric rhesus macaque (RM) models of HIV infection and treatment can be instrumental (19). Building on pediatric RM models of breast milk transmission (20) and persistence (21) with simian immunodeficiency viruses (SIVs), here we have developed a pediatric RM model of ART and viral rebound using infant RMs experimentally infected with a next generation chimeric simian-human immunodeficiency virus, SHIV.CH505.375H.dCT (22), that would permit an assessment of interventions directed against the HIV-Env. This virus carries a mutation in the CD4 binding site that facilitates entry via rhesus CD4 molecule that has been previously demonstrated to replicate efficiently in adult RMs (22), recapitulating the viral replication dynamics and immunopathogenesis of HIV infection in humans (23). We used the infant RM model to characterize the replication kinetics and virus-specific humoral immune responses during short-term ART and after ATI. We also utilized a unique opportunity to compare the viral and immune response kinetics of infant monkeys to that of adults infected with the same virus, during ART and post-ATI. Furthermore, we validated and assessed these RM models by examining clinically established biomarkers of time to viral rebound and explored the relationship between immune response and viral rebound. This infant RM ATI model will be a valuable addition to the HIV cure research toolbox to guide translational studies for evaluating the efficacy of therapeutic strategies towards attaining drug-free HIV remission for children.

## RESULTS

### Kinetics of SHIV.CH505.375H.dCT replication in orally infected infant RMs

Six infant RMs were orally challenged with SHIV.CH505.375H.dCT (22), as described in methods. Kinetics of SHIV replication in these infants were monitored for 12 weeks post infection (w.p.i), when they were initiated on a daily subcutaneous ART regimen of tenofovir disoproxil fumerate (TDF), emtricitabine (FTC), and dolutegravir (DTG) for 8 weeks. After 8 weeks of ART, treatment was interrupted and infants were monitored for an additional 8 weeks followed by necropsy (Figure 1A).

**Figure 1:**
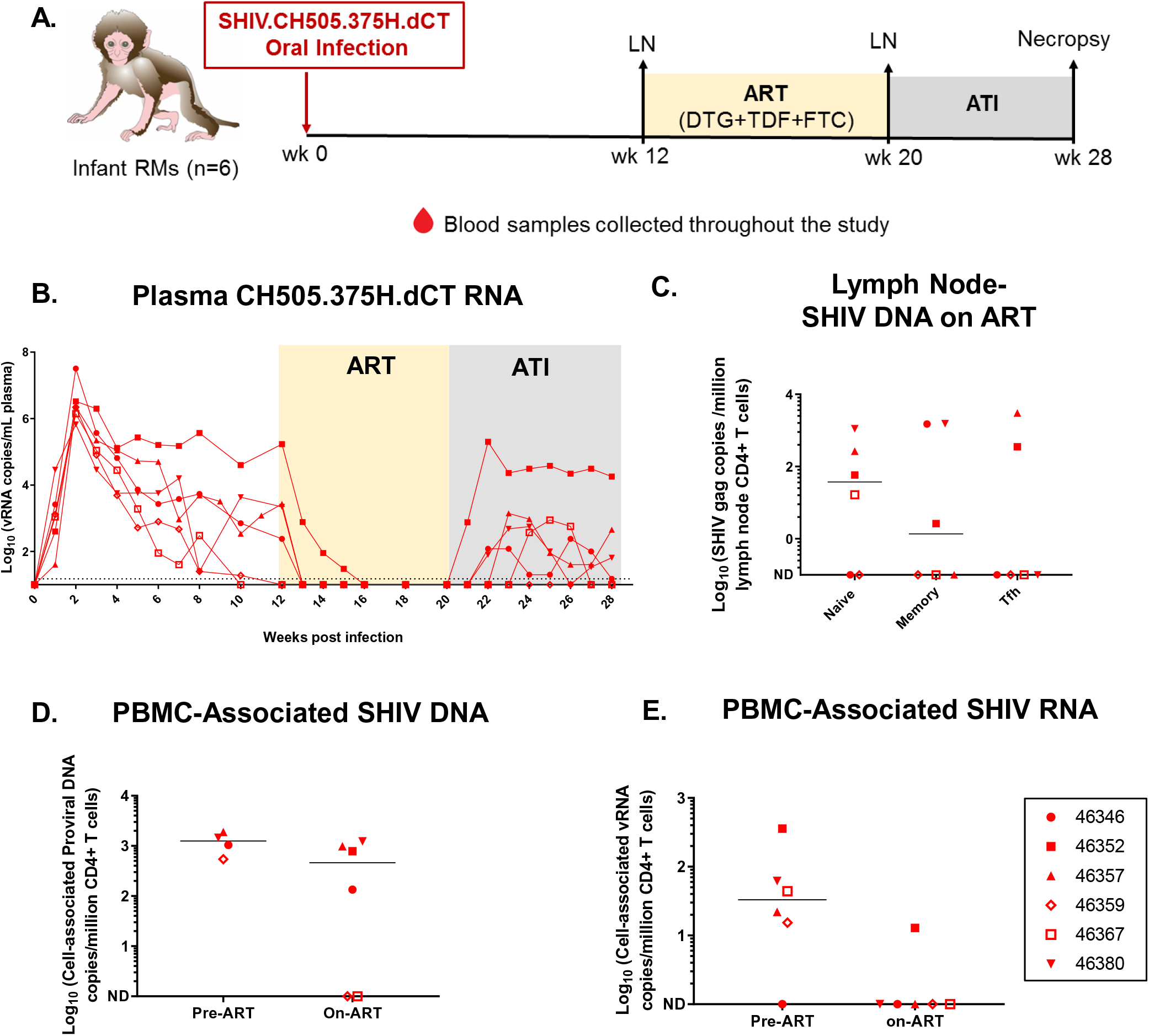
SHIV.CH505.375H.dCT replication kinetics prior to and following ATI in infant RMs. **(A)** Schematic representation of SHIV.CH505.375H.dCT infection (0-12 weeks), ART (12-20 weeks) and ATI (20-28 weeks) in infant RMs. Blood samples were collected at weekly intervals throughout the study and peripheral lymph nodes (LN) were collected at 12 w.p.i and on ART (20 w.p.i) **(B)** Kinetics of plasma SHIV RNA over 28 weeks was measured by qRT-PCR. **(C)** Peripheral lymph nodes from RMs on ART (20 w.p.i) were collected, and naïve, memory and Tfh CD4+ T cell-associated SHIV DNA was estimated by qPCR. **(D)** Cell-associated SHIV DNA (CA-SHIV DNA) and **(E)** cell-associated SHIV RNA (CA-SHIV RNA) per million CD4+ T cells in the peripheral blood was monitored by ddPCR in the infant RMs pre-ART (6 w.p.i) and on ART (18 w.p.i). The sensitivity of the ddPCR assay was detection of 1 SHIV gag copy in 10,000 CD4+ T cells. Therefore, only those animals that had ≥10,000 CD4+ T cells at a particular time point, were included in the analysis. Each symbol represents an individual animal. Yellow and grey boxes represent duration of ART (week 12-20) and duration of ATI (week 20-28), respectively. Medians are indicated as horizontal lines on the dot plots. Infants with plasma VL<15 copies/mL at 12 w.p.i has been represented with open symbols.

In the acute phase of infection, plasma viral load (VL) peaked at 2 w.p.i (6.7×10^5^-3.2×10^7^ vRNA copies/mL of plasma), and then declined over time (Figure 1B). Most of the RMs did not achieve a stable VL set point, and 2 had plasma VL< limit of detection (LOD) of 15 copies/mL before ART initiation (Table 1). Of note, 1 of these 2 infants was most resistant to infection (Table 1), and neither had an MHC allele previously associated with SHIV control (24, 25) (Table S1). CD4+ T cell frequencies were generally stable, with a slight decrease in the median frequency between 2-3 w.p.i. (Figure S1A), similar to the transient peripheral CD4+ T cells decline in acute HIV infection.

**Table 1.** SHIV.CH505.375H.dCT infected rhesus macaques, weeks of challenges to infection, age at infection, sex, time to viral control post-ART and time to viral rebound post-ATI.

| Animal Age | Animal ID | Challenges to infection (wk) | Age at infection<sup>a</sup> | Sex<sup>b</sup> | Pre- ART control<sup>c,e</sup> | Time to viral control post-ART (wk)<sup>d,f</sup> | Viral rebound post ATI<sup>c,g</sup> | Time to viral rebound post-ATI (wk)<sup>d</sup> | Post- rebound control<sup>c,d,h</sup> |
| --- | --- | --- | --- | --- | --- | --- | --- | --- | --- |
| Infant | 46357 | 1 | 5 | M | N | 1 | Y | 3 | N |
|  | 46346 | 2 | 9 | F | N | 1 | Y | 6 | Y |
|  | 46352 | 2 | 9 | F | N | 4 | Y | 1 | N |
|  | 46359 | 3 | 10 | F | Y | N/A | N | N/A | N/A |
|  | 46367 | 7 | 14 | M | Y | N/A | Y | 4 | Y |
|  | 46380 | 4 | 11 | F | N | 1 | Y | 3 | N |
| Adult | 39472 | 1 | 8 | F | N | 1 | N | N/A | N/A |
|  | 43068 | 1 | 5 | F | N | 1 | N | N/A | N/A |
|  | 43268 | 1 | 4 | F | N | 1 | Y | 3 | N |
|  | 42368 | 1 | 5 | F | N | 1 | Y | 3 | Y |
|  | 39950 | 1 | 8 | F | N | 4 | Y | 2 | N |
|  | 38200 | 1 | 10 | F | N | 1 | N | N/A | N/A |

<sup>a</sup>Weeks in infants; years in adults
<sup>b</sup>M, male; F, female
<sup>c</sup>N, No; Y, Yes
<sup>d</sup>N/A, Not applicable
<sup>e</sup>Pre ART control: plasma VL≤15 copies/mL at ART start (12 w.p.i)
<sup>f</sup>Viral control post ART: plasma VL≤15 copies/mL after ART start
<sup>g</sup>Viral rebound: Plasma VL≥10X detection limit of the assay (150 copies of vRNA/mL plasma) post ART discontinuation
<sup>h</sup>Post-rebound control: Plasma VL≤15 copies/mL after 8 weeks post-ATI, in monkeys showing viral rebound.

Upon ART initiation, the infant RMs demonstrated plasma VL<LOD within 1-4 weeks and none of them had detectable plasma VLs during the short course of ART. Upon ATI, 5 of 6 infants had viral rebound within 1-6 weeks (median: 3 weeks), of which 2 demonstrated plasma VL<LOD, within 2-3 weeks of viral rebound (Table 1). Interestingly, 1 of the 2 animals with VL< 15 copies prior to ART (46367), experienced viral rebound, post-ATI (Figure 1B). Not surprisingly, the animal with persistently high pre-ART viremia (46352) was the first to rebound post-ATI and experienced the highest rebound viremia.

### SHIV.CH505.375H.dCT reservoir in peripheral lymph nodes (LNs) and PBMCs of infant RMs

We assessed the size of viral reservoirs of infant RMs while they were virologically controlled on ART. SHIV DNA was quantified in peripheral LN-associated naïve, memory, and T follicular helper (Tfh) CD4+ T cells after 8 weeks of ART. Viral DNA was detected in all three CD4+ T cell populations in a subset of animals, with 4 of 6 monkeys having detectable DNA in naïve CD4+ T cells, 3 of 6 monkeys in memory CD4+ T cells, and 2 of 6 monkeys in Tfh cells (Figure 1C). Finally, we measured cell-associated SHIV-DNA (CA-SHIV DNA) and cell-associated SHIV-RNA (CA-SHIV RNA), per million CD4+ T cells isolated from PBMCs using digital droplet PCR (ddPCR). Of note, we could only report CA-SHIV DNA and CA-SHIV RNA data for those animals which had input cell counts greater than the threshold cell count for the assay (see methods). Our data demonstrated a decrease in CA-SHIV DNA (Figure 1D) and -SHIV RNA/million CD4+ T cells (Figure 1E), with only one infant (46352) having detectable CA-SHIV RNA, after 6 weeks of ART.

### Anatomic distribution of SHIV.CH505.375H.dCT post rebound in infant RMs

As anatomic sites of viral replication after ATI might reveal major sources of viral rebound, we sought to determine the distribution of SHIV.CH505.375H.dCT in blood and tissue compartments, at necropsy. Cell-associated infectious SHIV was measured in oral and gut-associated tissues (8 weeks post-ATI) using a Tzm-bl based co-culture assay (Figure S2), and 50% cellular infectious dose (CID_50_) for each tissue was reported (see methods). Our data demonstrated that the infectious virus was primarily distributed in the LNs and gut-associated tissues, compared to spleen (Figure 2A), which might be attributed to the lower proportion of CD4+ T cells in spleen (Figure S1B), vs. LNs. Interestingly, a higher number of animals had infectious virus detectable in oral LN (submandibular LN) compared to mesenteric LN and no cell-associated infectious virus was detected in PBMCs.

**Figure 2:**
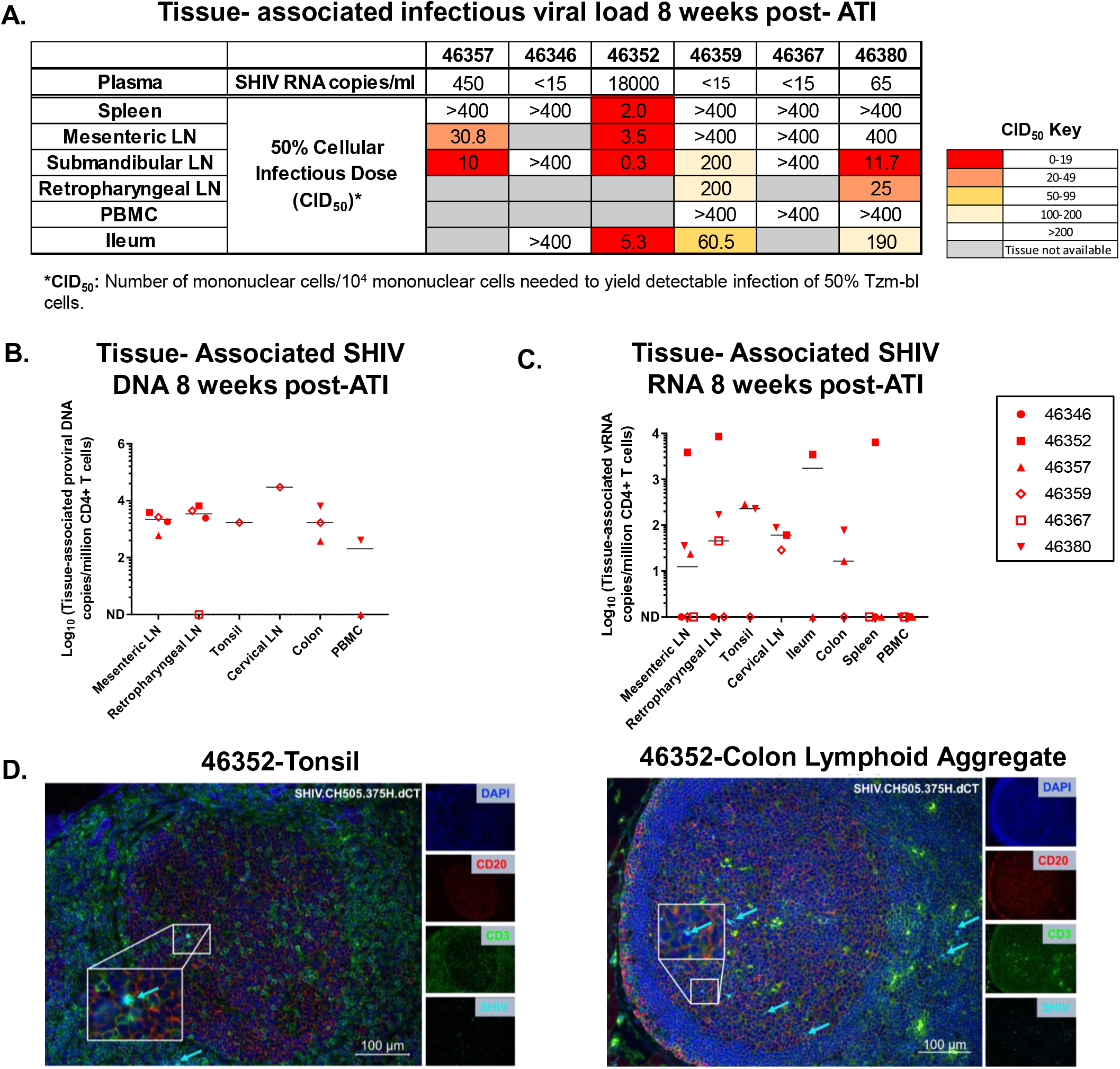
Tissue-associated infectious viral loads upon ATI, in mononuclear cells isolated from PBMCs, lymphoid and GI tissues of orally infected infant RMs. **(A)** Tissue-associated infectious SHIV.CH505.375H.dCT titers measured through tissue mononuclear cell coculture with Tzm-bl reporter cells. Reported titers represents the estimated minimum number of mononuclear cells per 10^4^ mononuclear cells required to yield detectable infection of 50% Tzm-bl cells (CID_50_). **(B)** CD4+ T cell-associated proviral DNA and **(C)** CD4+ T cell-associated viral RNA loads at necropsy (28 w.p.i) reported as copy number/million CD4+ T cells in PBMCs, lymphoid and GI tissue mononuclear cells. Each symbol represents one individual animal. Medians are indicated as horizontal lines on the dot plots. Infants with plasma VL<15 copies/mL at 12 w.p.i has been represented with open symbols. The sensitivity of the ddPCR assay was detection of 1 SHIV gag copy in 10,000 CD4+ T cells. Therefore, only those animals that had ≥10,000 CD4+ T cells at a particular time point, were included in the analysis. **(D)** Tonsil and colon sections from the SHIV.CH505.375H.dCT infected infant RM that demonstrated highest peak plasma VL post-rebound (20,000 vRNA copies/mL plasma). Tissue sections were stained with the nuclear marker DAPI (dark blue) to identify cells, and with antibodies specific for CD3 (green) and CD20 (red). Virally-infected cells were identified by *in-situ* hybridization (cyan). To better visualize the virally-infected cells, we magnified a specific region (white box) in each image. Each panel consists of a larger image with the overlay of all markers and 4 smaller side panels of the same field for each individual channel. Arrow colors correspond to the indicated marker. The large image has a scale bar in the lower right corner.

Eight weeks post-ATI, tissue-associated SHIV DNA and -RNA per million CD4+ T cells were detectable at variable levels (SHIV DNA: 1-3×10^4^ copies/million CD4+ T cells and SHIV RNA: 1-8.61×10^3^ copies/million CD4+ T cells) (Figure 2B and 2C). None of the monkeys had detectable SHIV RNA in PBMCs, further confirming our co-culture-based tissue-associated infectious viral load data. We further defined the anatomic distribution of CD3+SHIV+ cells in LN and gut-associated lymphoid tissues (GALT) of the infant that showed the highest plasma VL, post rebound, using a dual immunohistochemistry (IHC)/ in situ hybridization (ISH) approach (Figure 2D). Interestingly, CD3+SHIV+ cells were detected within the B cell follicles in addition to the T cell zone, suggesting that resident Tfh cells in both the tonsil and GALT can support viral replication.

### Adult RMs have comparable SHIV.CH505.375H.dCT replication kinetics, viral reservoir, and rebound-virus distribution compared to infants

We took the opportunity to compare the viral replication kinetics and reservoir in infant RMs to that of adult RMs infected with the same SHIV strain from a separate study, and thus a cohort of convenience (26). Six adult RMs, were intravenously infected with SHIV.CH505.375H.dCT (see methods) and started on triple ART regimen of TDF, FTC and DTG at 12 w.p.i. After 12 weeks of ART, therapy was discontinued, and the animals were euthanized 8 weeks post-ATI (Figure 3A). Similar to infant RMs, plasma VL in adults peaked at 2 w.p.i (3×10^5^-1.2×10^7^ copies of vRNA/mL of plasma). Additionally, the overall kinetics of plasma VL during acute SHIV infection was highly comparable between the two groups (Figure 3B). Upon ART initiation, plasma VL in adults was below LOD (<15 copies/mL plasma) within 1-4 weeks, with one monkey experiencing a viral blip (>15 copies/mL plasma), during the course of ART. Even though the ART regimen in adults was slightly longer than infants, 3 of 6 adults showed viral rebound within 2-3 weeks post-ATI. Interestingly, 1 of 3 adults that experienced viral rebound demonstrated plasma VL below LOD at 8 weeks post-ATI (Table 1).

**Figure 3:**
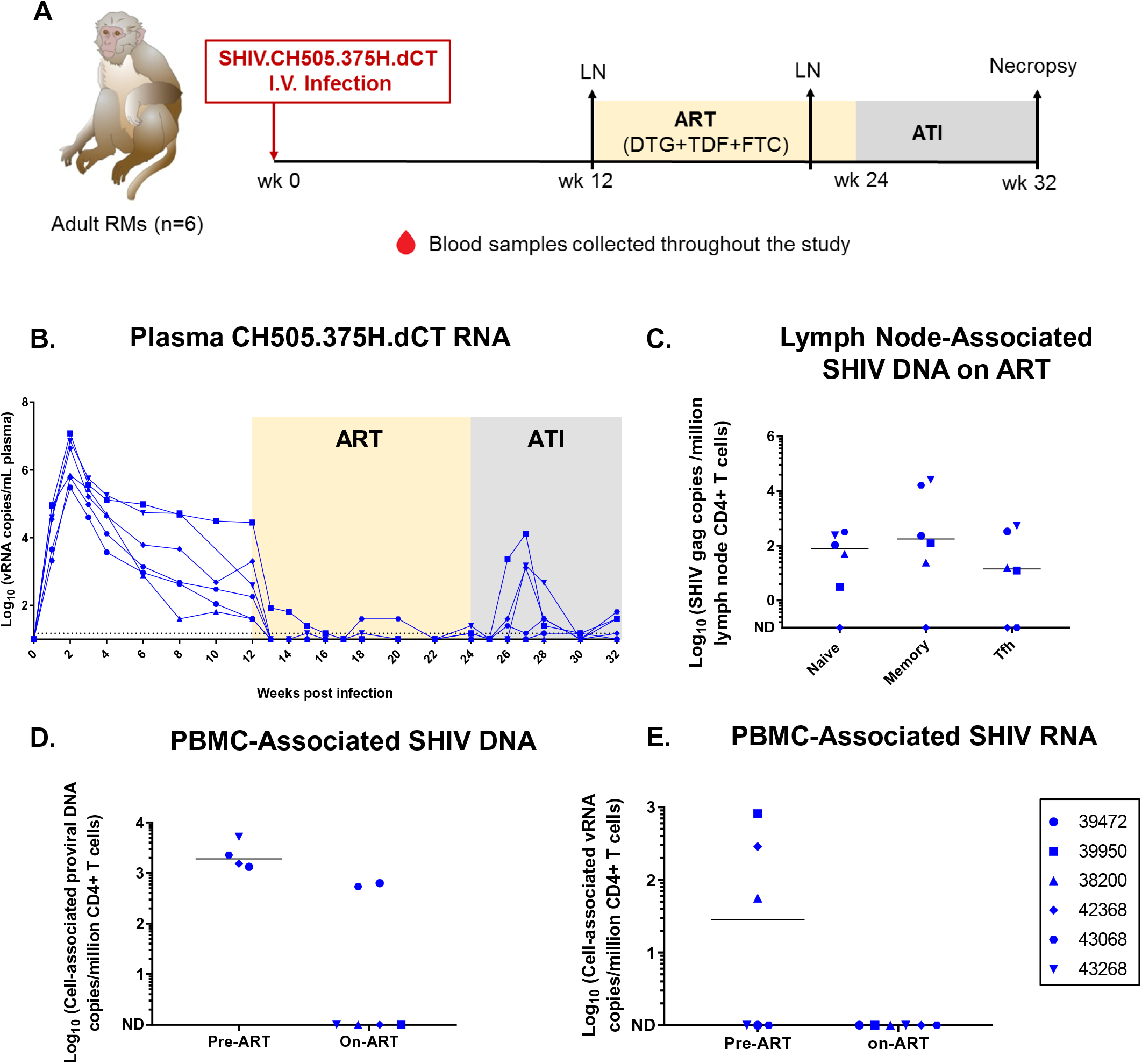
SHIV.CH505.375H.dCT replication kinetics prior to and following ATI in adult RMs. **(A)** Schematic representation of SHIV.CH505.375H.dCT infection (0-12 weeks), ART (12-24 weeks) and ATI (24-32 weeks) in adult RMs. Blood samples were collected at weekly intervals throughout the study and peripheral lymph nodes (LN) were collected at 12 w.p.i and after 8 weeks of ART (20 w.p.i) **(B)** Kinetics of plasma SHIV RNA over 32 weeks was measured by qRT-PCR. **(C)** Peripheral lymph nodes from macaques on ART (20 w.p.i) were collected, and naïve, memory and Tfh CD4+ T cell-associated SHIV DNA was estimated by qPCR. **(D)** Cell-associated SHIV DNA (CA-SHIV DNA) and **(E)** Cell-associated SHIV RNA (CA-SHIV RNA) from CD4+ T cells of peripheral blood pre-ART (6 w.p.i for DNA and 12 w.p.i for RNA) and on-ART (18 w.p.i) was monitored by ddPCR. The sensitivity of the ddPCR assay was detection of 1 SHIV gag copy in 10,000 CD4+ T cells. Therefore, only those animals that had ≥10,000 CD4+ T cells at a particular time point, were included in the analysis. Each symbol represents an individual animal. Yellow and grey boxes represent duration of ART (week 12-24) and duration of ATI (week 24-32), respectively. Medians are indicated as horizontal lines on the dot plots.

SHIV reservoir was detectable in adults in naïve, memory, and Tfh CD4+ T cell subsets of peripheral LNs. (Figure 3C). As observed with infant RMs, CA-SHIV DNA (Figure 3D) and CA-SHIV RNA/million CD4+ T cells (Figure 3E) declined upon ART, with only 2 adults demonstrating detectable CA-SHIV DNA levels after 6 weeks on ART. Like infant RMs, co-culture assays detected higher infectious viral titers in oral LN (submandibular) compared to mesenteric LN, whereas cell-associated infectious virus was not detected in PBMCs (Figure 4A). CA-SHIV DNA and RNA were detected in tissues at variable levels, with none of the animals having detectable CA-SHIV RNA in PBMCs (Figure 4B and 4C). Similar to infants, the tonsils and colon lymphoid aggregates of the adult RM that showed highest plasma VL post-rebound had detectable CD3+SHIV+ cells within the B cell follicle, in addition to T cell zone at necropsy (Figure 4D).

**Figure 4:**
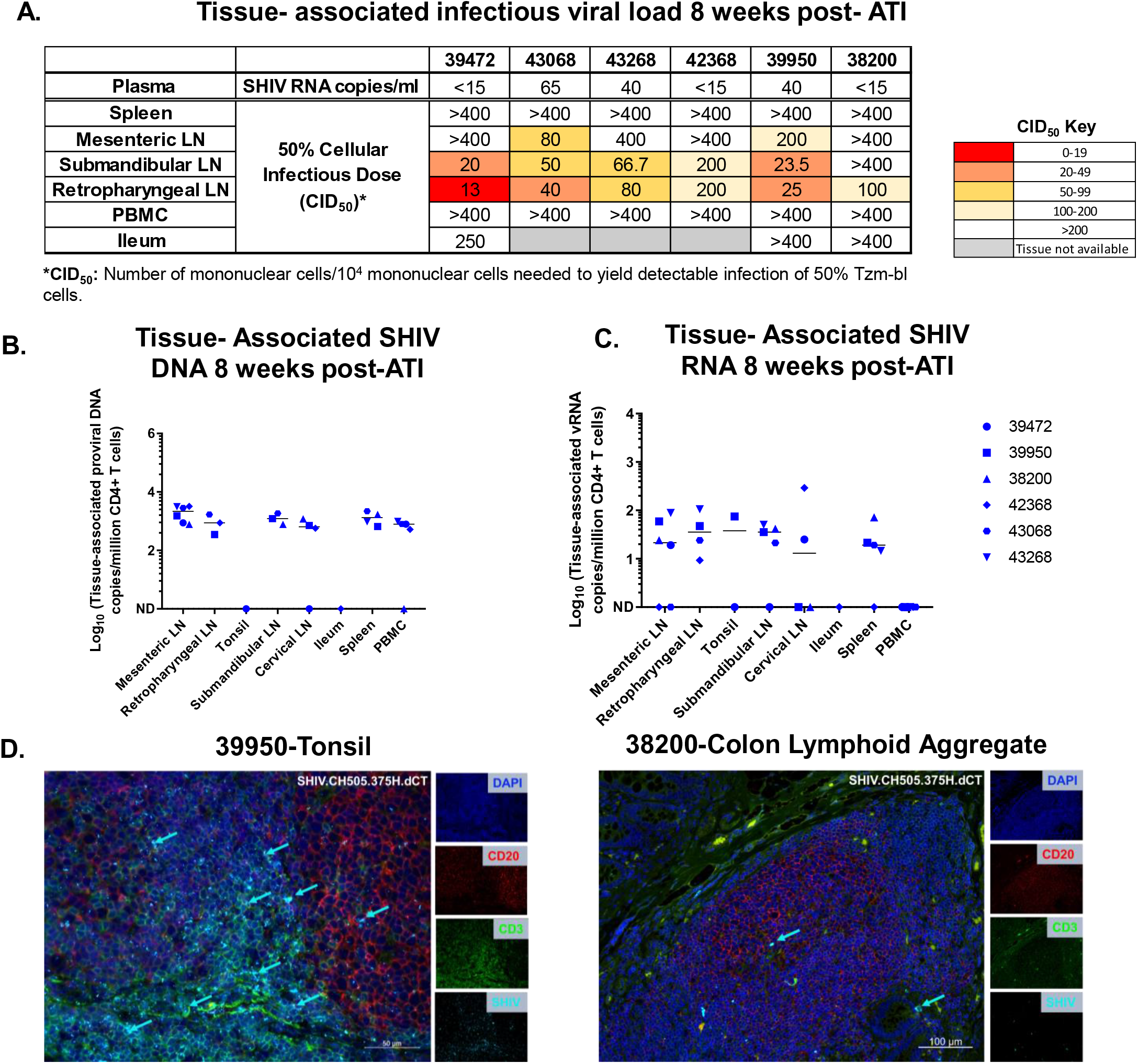
Tissue-associated infectious virus load upon ATI, in mononuclear cells isolated from PBMCs, lymphoid and GI tissues of adult RMs intravenously infected with SHIV.CH505.375H.dCT. **(A)** Tissue-associated infectious SHIV.CH505.375H.dCT titers measured through tissue mononuclear cell coculture with Tzm-bl reporter cells. Reported titers represents the estimated minimum number of mononuclear cells per 10^4^ mononuclear cells required to yield detectable infection of 50% Tzm-bl cells (CID_50_). **(B)** CD4+ T cell-associated proviral DNA and **(C)** Viral RNA loads reported as copy number/million CD4+ T cells in PBMCs, lymphoid and GI tissue mononuclear cells. Each symbol represents one individual monkey at necropsy (week 32 pi). Medians are indicated as horizontal lines on the dot plots. **(D)** Tonsil and colon sections from the SHIV.CH505.375H.dCT infected adult RM (39950) that demonstrated highest peak plasma VL post-rebound (13000 vRNA copies/mL plasma). Tissue sections were stained with the nuclear marker DAPI (dark blue) to identify cells, and with antibodies specific for CD3 (green) and CD20 (red). Virally-infected cells were identified by *in-situ* hybridization (cyan). Each panel consists of a larger image with the overlay of all markers and 4 smaller side panels of the same field for each individual channel. Arrow colors correspond to the indicated marker. The large image has a scale bar in the lower right corner.

### Viral diversity before ART initiation and post-ATI in infant and adult RMs

As the viral kinetics during acute infection, ART, and post-ATI were similar in infant and adult monkeys, we next sought to compare plasma viral Env diversity between the infant (46352) and adult RM (39950) that showed highest plasma VL post-rebound. We performed single genome amplification (SGA) and sequencing of the *env* gene pre-ART and post-ATI from plasma and calculated average pairwise distance (APD) within Env sequences. In the infant 46352, pre-ART (12 w.p.i) viral Env was more homogeneous (average pairwise distance: APD: 0.0006), compared to 2 weeks (APD: 0.0018) or 8 weeks post-ATI viral Env (APD:0.0038) (Figure 5A). In contrast, for the adult 39950, pre-ART viral Env (12 w.p.i) had a 4-fold higher diversity (APD: 0.004), compared to 2 weeks post-ATI viral Env (APD: 0.001) (Figure 5B).

**Figure 5.**
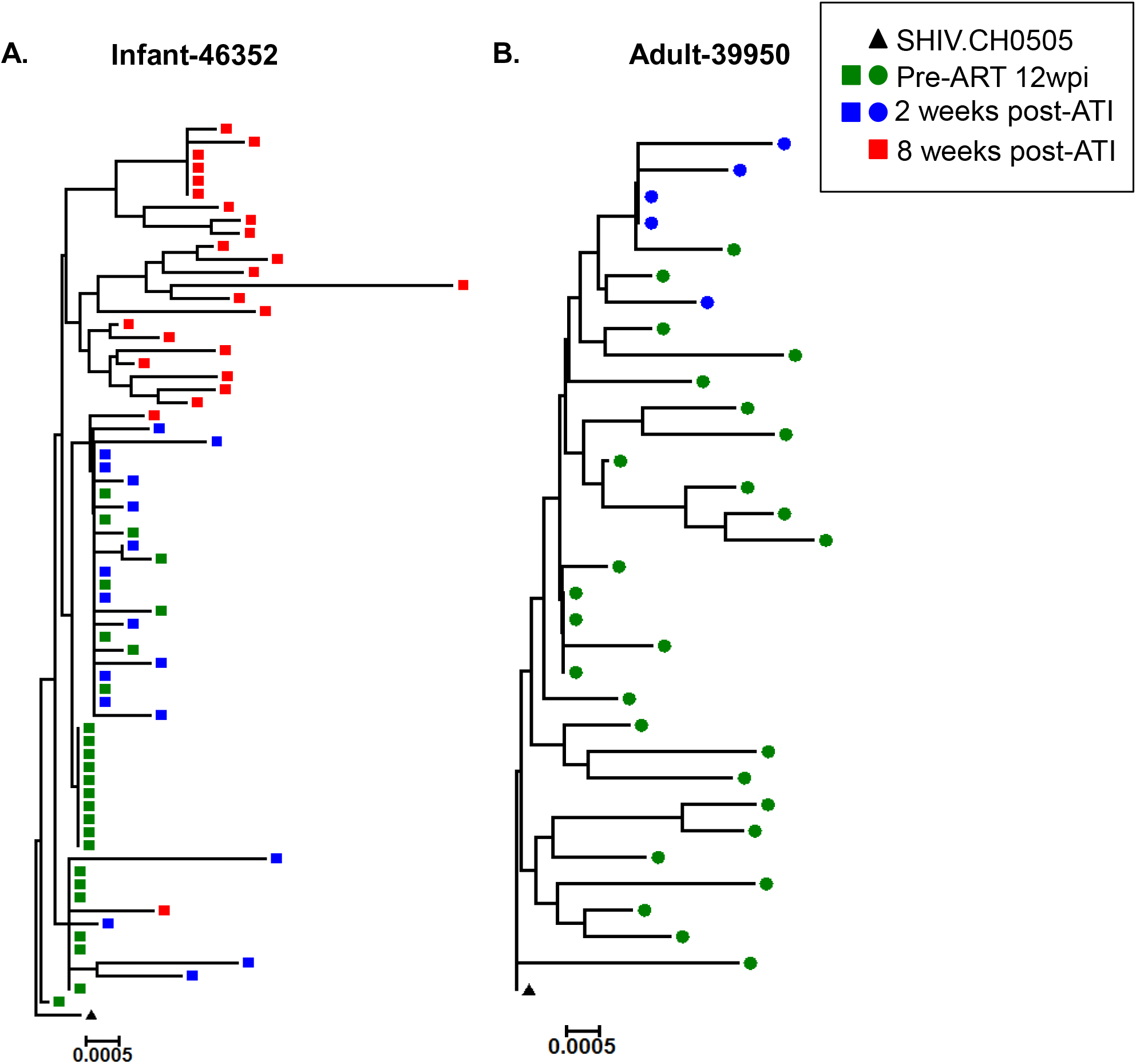
Phylogenetic tree analysis of the *env* gene sequences obtained from pre-ART and post-ATI plasma of infant and adult RM demonstrating highest peak plasma VL post-rebound. Standard SGA techniques were used to analyze the *env* gene from pre-ART and post-ATI samples of infants and adult RMs with highest peak rebound plasma VL. Phylogenetic tree representing viral Env diversity in **(A)** infant 46352 and **(B)** adult 39950.

### ART dampens the magnitude of humoral responses in SHIV-infected infant and adult RMs

As previous reports have noted a loss of HIV-specific humoral responses in human infants on ART (27), we investigated differences in the kinetics, magnitude, and breadth of the Env-specific humoral responses on ART and following ATI between the age groups. In both age groups, all monkeys developed detectable autologous gp120-specfic IgG responses at 12 w.p.i. (Figure 6A). After 8 weeks of ART, gp120-specific IgG response declined in both groups, the decline being more pronounced in infants. Yet, the gp120-specific IgG response rebounded in both groups post-ATI. Interestingly, the infant (46359) with plasma VL<15 copies pre-ART and no viral rebound, developed similar gp120-specific IgG response to that of other infants. We then mapped the Env-domain specificity of the antibody responses, and observed dominant responses against the V3 and -C5 linear epitopes in both groups (Figure S3), which was not altered upon ART. Interestingly, ART initiation completely abrogated plasma antibodies against the CD4 binding site and only 2 of 6 infants and no adults regained this response within the 8 weeks of follow-up after ART was discontinued. (Figure 6B).

**Figure 6:**
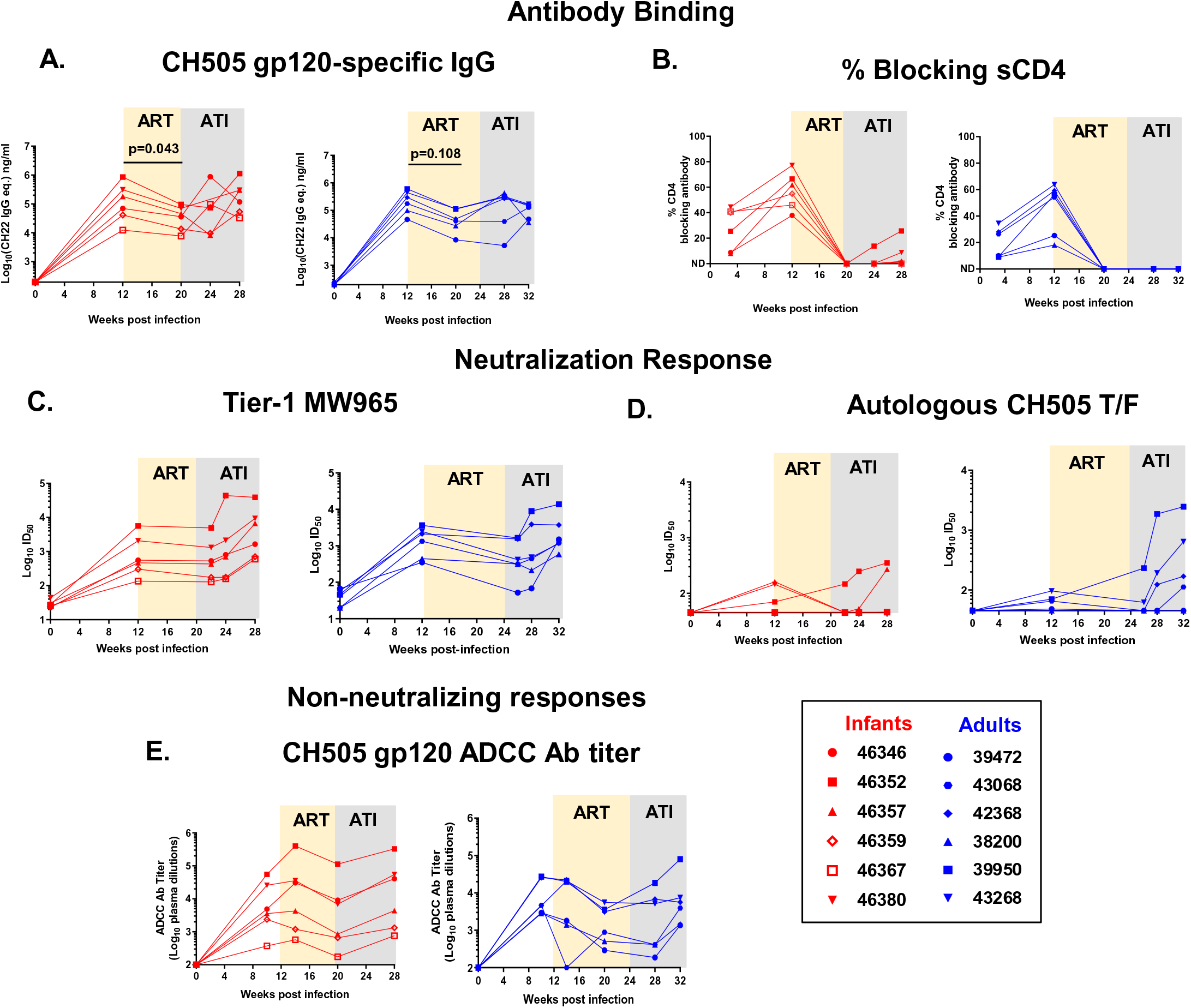
Magnitude and kinetics of humoral responses to acute SHIV.CH505.375H.dCT infection, during ART and ATI in RMs. Plasma from infant and adult rhesus macaques were analyzed for. **(A)** HIV CH505 gp120 IgG response **(B)** blocking of soluble CD4-gp120 interactions **(C)** tier-1 neutralization response against MW965 **(D)** neutralization response against CH505 T/F, and **(E)** ADCC titer against CH505 gp120-coated target cells. Red symbols represent infant RMs and blue symbols represent adult RMs. Each symbol represents an individual macaque. Yellow and grey boxes represent duration of ART and duration of ATI in the RMs, respectively. Infants with plasma VL<15 copies/mL at 12 w.p.i has been represented with open symbols. P values were calculated using Wilcoxon signed rank test.

We next evaluated the HIV neutralization potency of RM plasma against a tier-1 clade-matched isolate MW965 and the autologous tier-2 CH505 virus. Both infants and adults developed neutralization activity against MW965 by 12 w.p.i, which continued to increase post-ATI, with equal potency between age groups (Figure 6C). However, only half of both the adult and infant RMs had a detectable neutralization response against autologous CH505 at 12 w.p.i. (Figure 6D). Post-ATI, 2 of 6 infants and 4 of 6 adults demonstrated increasing neutralization responses against the autologous virus. In both age groups, gp120-specific ADCC titers dampened upon ART initiation, yet recovered to pre-ART levels after ATI (Figure 6E).

### T cell activation during ART in infant and adult RMs

Systemic immune activation has been associated with HIV replication and poor disease outcomes (28). Furthermore, T cell exhaustion markers have been reported as predictors of viral rebound in a human study (15). Therefore, we assessed the activation and exhaustion status of T cells from SHIV-infected, infant and adult RMs. Unlike adults, infant-activated, -proliferating and - exhausted CD4+ T cell populations increased at 10 w.p.i compared to pre-infection levels, which might be attributed to the age-specific development of T cell populations (Figure 7A and 7B).

**Figure 7:**
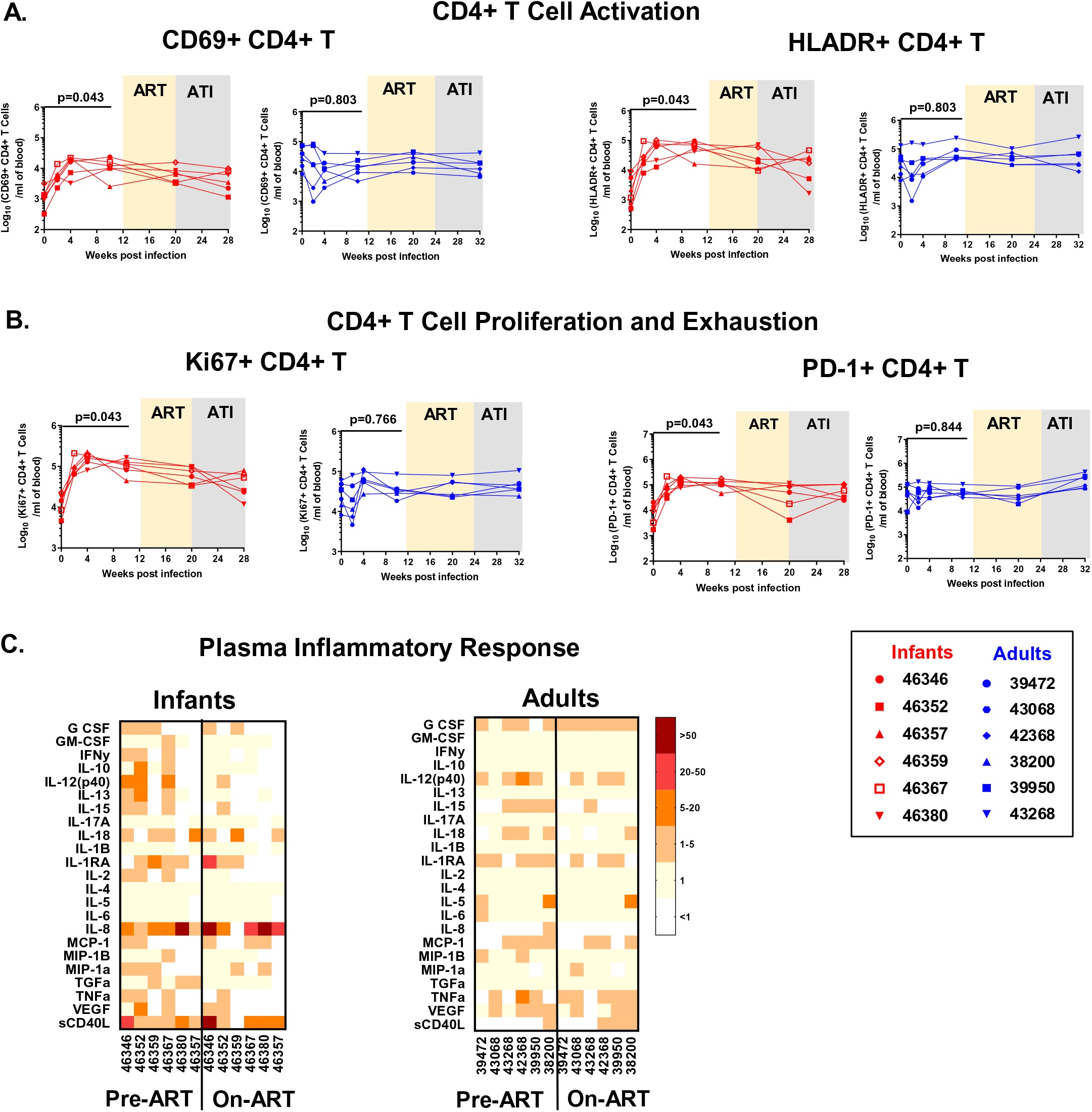
CD4+ T cell activation and plasma inflammatory responses in SHIV.CH505.375H.dCT infected infant and adult RMs. Absolute counts per ml blood of **(A)** activated (CD69+, HLADR+) CD4+ T cells and **(B)** proliferating (Ki67+ CD4+) and exhausted (PD-1+) CD4+ T cells. Red symbol represents infant RMs and blue symbols represents adult RMs. Each symbol represents one animal. Yellow and grey boxes represent duration of ART and duration of ATI, respectively. **(C)** Plasma from infected infant and adult RMs were analyzed by multiplexed luminex assay for expression of cytokines before ART (week 12) and 8 weeks on ART (week 20). Heat map represents fold change of each analyte over pre-infection plasma levels. Infants with plasma VL<15 copies/mL at 12 w.p.i has been represented with open symbols. P values were calculated using Wilcoxon signed rank test.

Furthermore, we measured the concentrations of inflammatory chemokines and cytokines in RM plasma pre-ART (12 w.p.i) and on ART (8 weeks of ART). While infant plasma demonstrated a higher magnitude and breadth of cytokine and chemokine levels than adults, no notable difference was observed pre-ART vs. on ART in both age groups (Figure 7C).

### Correlates of time to SHIV rebound in infant and adult RMs

To validate our established RM SHIV rebound models in identifying correlates of time to viral rebound, we performed a univariate cox-proportional hazard modeling on a subset of the measured virologic and immunologic parameters, after adjustment for age. We defined time to viral rebound as a plasma VL >10 times LOD (150 copies/mL) post-ART discontinuation. Plasma VL pre-ART, ADCC antibodies, CD4+ CD69+ T cells and autologous virus neutralizing antibodies were pre-selected as primary parameters based on their previously published associations with rate of HIV acquisition (29, 30) and disease progression (31, 32). Of these parameters, higher levels of plasma VL pre-ART and ADCC antibody titers on ART demonstrated associations with increased risk of viral rebound (Table 2). Next, we applied the model on an additional set of virologic and immunologic parameters. The analysis identified higher levels of CH505 gp120-specific IgG responses pre-ART and on ART, tier-1 MW965 neutralizing antibody titer, and %sCD4 blocking antibodies pre-ART to be associated with higher risk of rebound (Table 2), suggesting that our macaque models were suitable for monitoring correlates of viral rebound.

**Table 2.** Virologic and immunologic correlates of time to viral rebound, with adjustment for age.

| Variables | HR (95% CI) | Unadjusted p value | Adjusted p value |
| --- | --- | --- | --- |
| **Pre-selected Variables** |  |  |  |
| Plasma VL at ART start<sup>a</sup> | 2.2 (1.24-3.95) | **0.008** | NA |
| ADCC antibody titer on ART<sup>a</sup> | 1.9 (1.05-3.45) | **0.034** | NA |
| CD4+ CD69+ T cells on ART<sup>b</sup> | 0.82 (0.26-2.56) | 0.727 | NA |
| CH505 T/F neutralizing antibody titer at ART start<sup>a</sup> | 2.1 (0.82-5.57) | 0.119 | NA |
| **Exploratory Variables** |  |  |  |
| **(a) Virologic and humoral responses** |  |  |  |
| Peak plasma VL<sup>a</sup> | 1.39 (0.86-2.27) | 0.182 | 0.364 |
| CH505 gp120-specific IgG pre ART<sup>a</sup> | 4.61 (1.38-15.42) | **0.0131** | 0.066 |
| CH505 gp120-specific IgG on ART<sup>a</sup> | 6.06 (1.33-27.54) | **0.0197** | 0.066 |
| Tier-1 MW965 neutralizing antibody pre-ART<sup>a</sup> | 4.12 (1.42-11.96) | **0.00926** | 0.066 |
| sCD4 blocking antibodies pre-ART<sup>c</sup> | 1.07 (1-1.13) | **0.0402** | 0.1 |
| **(b) CD4+ T cell phenotypes and inflammatory responses** |  |  |  |
| CD4+ T pre-ART<sup>b</sup> | 0.97 (0.86-1.1) | 0.66 | 0.81 |
| CD4+ T on ART<sup>b</sup> | 0.99 (0.9-1.09) | 0.77 | 0.82 |
| HLADR+ CD4+ T on ART<sup>b</sup> | 1 (1-1) | 0.71 | 0.81 |
| Ki67+ CD4+ T on ART<sup>b</sup> | 1 (0.98-1.05) | 0.4 | 0.8 |
| PD-1+ CD4+ T on ART<sup>b</sup> | 1 (1-1) | 0.32 | 0.8 |
| G-CSF on ART<sup>b</sup> | 1.1 (0.91-1.28) | 0.36 | 0.8 |
| IL-12 on ART<sup>d</sup> | 1.1 (0.96-1.36) | 0.13 | 0.8 |
| IL-15 on ART<sup>d</sup> | 0.77 (0.4-1.5) | 0.45 | 0.8 |
| IL-18 on ART<sup>d</sup> | 0.97 (0.93-1.01) | 0.15 | 0.8 |
| IL-1RA on ART<sup>d</sup> | 1 (0.99-1.01) | 0.71 | 0.81 |
| IL-8 on ART<sup>d</sup> | 1 (1-1) | 0.97 | 0.97 |
| MCP-1 on ART<sup>d</sup> | 1 (0.99-1.02) | 0.34 | 0.8 |
| MIP1a on ART<sup>d</sup> | 0.68 (0.3-1.57) | 0.37 | 0.8 |
| TNFa on ART<sup>d</sup> | 0.91 (0.79-1.04) | 0.17 | 0.8 |
| VEGF on ART<sup>d</sup> | 0.97 (0.86-1.1) | 0.64 | 0.81 |
| SCD40L on ART<sup>d</sup> | 1 (1-1) | 0.7 | 0.81 |

Values expressed as <sup>a</sup>Log<sub>10</sub>; <sup>b</sup>Absolute counts/nL of blood; <sup>c</sup>Percentages; <sup>d</sup>Concentrations
HR: Hazard ratio; CI: Confidence Interval; NA: Not applicable since multiple adjustment was not performed on pre-selected parameters.

Finally, we wanted to determine the differential impact of age on the correlates of viral rebound. Due to the relatively small sample size of the two age groups, we performed Kendall’s Tau rank correlation of each of the experimental parameters with viral rebound. In infants, lower plasma VL at ART start was associated with longer time to viral rebound and a correlation trend for longer time to viral rebound was observed for infant RMs with lower CH505 gp120-specific IgG responses pre-ART and on ART. In adults, lower values of plasma VL at ART start, peak plasma VL, CH505 gp120-specific IgG response pre-ART and Tier-1 MW965 neutralizing antibody titer at ART start were associated with longer time to rebound. Additionally, a correlation trend for longer time to viral rebound was observed for adult monkeys with lower CH505 gp120-specific IgG responses on ART (Table 3).

**Table 3.** Virologic and immunologic correlates of time to viral rebound in each age group.

| Variables | Infant |  |  | Adult |  |  |
| --- | --- | --- | --- | --- | --- | --- |
|  | Kendall's Tau | Unadjusted p value | Adjusted p value | Kendall's Tau | Unadjusted p value | Adjusted p value |
| **Pre-selected Variables** |  |  |  |  |  |  |
| Plasma VL at ART start<sup>a</sup> | -0.786 | **0.032** | NA | -0.886 | **0.022** | NA |
| ADCC antibody titer on ART<sup>a</sup> | -0.414 | 0.251 | NA | -0.701 | 0.064 | NA |
| CD4+ CD69+ T cells on ART<sup>b</sup> | 0.276 | 0.444 | NA | -0.078 | 0.837 | NA |
| CH505 T/F neutralizing antibody titer at ART start<sup>a</sup> | -0.54 | 0.152 | NA | -0.322 | 0.404 | NA |
| **Exploratory Variables** |  |  |  |  |  |  |
| **(a) Virologic and humoral responses** |  |  |  |  |  |  |
| Peak plasma VL<sup>a</sup> | 0 | 1 | 1 | -0.856 | **0.024** | **0.04** |
| CH505 gp120-specific IgG pre ART<sup>a</sup> | -0.69 | **0.056** | 0.14 | -0.856 | **0.024** | **0.04** |
| CH505 gp120-specific IgG on ART<sup>a</sup> | -0.69 | **0.056** | 0.14 | -0.701 | 0.064 | 0.08 |
| Tier-1 MW965 neutralizing antibody pre-ART<sup>a</sup> | -0.552 | 0.126 | 0.158 | -0.856 | **0.024** | **0.04** |
| sCD4 blocking antibodies pre-ART<sup>c</sup> | -0.552 | 0.126 | 0.158 | -0.545 | 0.15 | 0.15 |
| **(b) CD4+ T cell phenotypes and inflammatory responses** |  |  |  |  |  |  |
| CD4+ T pre-ART<sup>b</sup> | 0.138 | 0.702 | 0.936 | 0.078 | 0.837 | 0.837 |
| CD4+ T on ART<sup>b</sup> | 0 | 1 | 1 | 0.234 | 0.537 | 0.781 |
| HLADR+ CD4+ T on ART<sup>b</sup> | 0.276 | 0.444 | 0.936 | -0.078 | 0.837 | 0.837 |
| Ki67+ CD4+ T on ART<sup>b</sup> | -0.276 | 0.444 | 0.936 | 0.078 | 0.837 | 0.837 |
| PD-1+ CD4+ T on ART<sup>b</sup> | 0.138 | 0.702 | 0.936 | 0.545 | 0.15 | 0.781 |
| G-CSF on ART<sup>b</sup> | 0.357 | 0.33 | 0.936 | -0.418 | 0.289 | 0.781 |
| IL-12 on ART<sup>d</sup> | -0.357 | 0.33 | 0.936 | -0.261 | 0.511 | 0.781 |
| IL-15 on ART<sup>d</sup> | -0.138 | 0.702 | 0.936 | 0.322 | 0.404 | 0.781 |
| IL-18 on ART<sup>d</sup> | 0.552 | 0.126 | 0.936 | 0.405 | 0.343 | 0.781 |
| IL-1RA on ART<sup>d</sup> | 0.138 | 0.702 | 0.936 | 0.389 | 0.304 | 0.781 |
| IL-8 on ART<sup>d</sup> | 0.138 | 0.702 | 0.936 | 0.078 | 0.837 | 0.837 |
| MCP-1 on ART<sup>d</sup> | -0.414 | 0.251 | 0.936 | -0.234 | 0.537 | 0.781 |
| MIP1a on ART<sup>d</sup> | 0.077 | 0.838 | 0.967 | 0.405 | 0.343 | 0.781 |
| TNFa on ART<sup>d</sup> | 0.357 | 0.33 | 0.936 | 0.234 | 0.537 | 0.781 |
| VEGF on ART<sup>d</sup> | -0.071 | 0.846 | 0.967 | 0.087 | 0.827 | 0.837 |
| SCD40L on ART<sup>d</sup> | 0 | 1 | 1 | -0.234 | 0.537 | 0.781 |

Values expressed as <sup>a</sup>Log<sub>10</sub>; <sup>b</sup>Absolute counts/nL of blood; <sup>c</sup>Percentages; <sup>d</sup>Concentrations
NA: Not applicable since multiple adjustment was not performed on pre-selected parameters.

## DISCUSSION

Identification of plasma viral RNA (31) and CD4+ T cell counts (33) as surrogate markers of HIV disease progression were instrumental in the development of ART for attaining improved clinical care in HIV-infected patients. As the HIV field rapidly turns towards achieving functional “cure”, there has been renewed interest in identifying biomarkers of viral rebound. Monitoring biomarkers will guide clinical trials with effective therapeutic candidates, minimizing investment to those that are unlikely to result in delay of viral rebound. However, ATI trials for identification of biomarkers remain logistically challenging, particularly in children, necessitating the development of tractable pediatric animal models. While pediatric ATI models of SIV-infected RMs are not new to the HIV field (34), clinically translatable RM models of ATI using SHIVs have been advancing (35). Thus, we have sought to establish an oral SHIV-infected infant ATI model with a previously validated ART regimen (21).

This study investigated the impact of ART on SHIV replication and SHIV-specific immune responses in the context of the maturing infant immune system. Employing an adult RM SHIV-infected cohort of convenience, we also demonstrated that infant and adult RMs have a comparable SHIV replication kinetics and are equally equipped in mounting immune responses, both on ART and post-ATI. Furthermore, we validated our RM ATI models by confirming previously reported virologic correlates of rebound and identified additional potential humoral immune correlates.

In this study, the infant and adult RMs were infected with a clade C SHIV variant, SHIV.CH505.375H.dCT, due to the predominance of clade C viruses in sub-Saharan Africa, where most pediatric HIV infections occur (36). In both infant and adult models, this SHIV variant achieved a peak viral load comparable to previously described SIV/SHIV models (21, 37), yet did not achieve a viral “set point” in most of the monkeys (Figure 1B and 3B). Additionally, a few monkeys from our cohort achieved natural virologic control, which is not uncommon in previously described SHIV infection models (38, 39). Moreover, our study revealed a comparable plasma VL and rebound kinetics in the two age groups.

Previous studies have claimed that infants may have impaired immune responses to HIV infection compared to adults (40). However, we previously demonstrated that vaccination of infants can induce robust HIV Env-specific IgG responses (41). Moreover, we recently reported that infant monkeys are capable of mounting durable anti-HIV humoral immune responses during acute SHIV infection, despite their maturing immune landscape (26). In this study, we observed comparable humoral immune responses between the two age groups during ART and post-ATI (Figure 6). Interestingly, a decline of HIV gp120-specific IgG responses and ADCC responses were observed on ART, without a change in the specificities of the Env domain-specific antibodies (Figure S3). A similar observation was made in human infants, where HIV-specific antibody levels decreased as the duration of ART increased (42), suggesting that circulating HIV antigen is a major driving factor for production of HIV Env-specific antibodies. In fact, plasma Env-specific IgG levels could provide a more comprehensive measure of viral replication in tissue sanctuaries which might not be reflected in the plasma viral load.

To identify correlates of viral rebound in infants and adults, we analyzed a comprehensive panel of 25 virologic and immunologic parameters. Our data confirmed a key, well-established clinical virologic marker, pre-ART plasma VL, as a correlate of viral rebound in both the age groups. Additionally, higher peak plasma VL was identified as a correlate of quicker viral rebound in adults, but not infants. This association of pre-therapy plasma VL with viral rebound has been demonstrated previously in an HIV gag-based therapeutic trial, where lower pre-ART plasma VL was independently associated with a lower post-ATI plasma VL (43). Among the immunologic parameters tested, in adults, higher gp120-specific IgG responses and neutralizing antibodies against a tier-1 virus correlated with quicker viral rebound, whereas in infants higher gp120-specific IgG responses, but not neutralizing antibodies against a tier-1 virus showed a correlation trend with quicker viral rebound. There has been continued interest in considering anti-HIV gp120 responses as a screening marker for ongoing viral replication or breakthrough, during suppressive ART (44). Additionally, heterologous neutralizing antibody responses at the time of treatment interruption has been associated with reduced viral load over time (45). However, these identified humoral responses have not been previously associated with time to viral rebound, and therefore, should be examined in future long-term studies. In resource limited settings, monitoring HIV-specific humoral responses in infants on suppressive ART might be beneficial due to small sample volume requirement and relatively low cost and technology burden compared to nucleic acid-based assays or quantitative viral outgrowth assays (QVOA).

There were a few notable limitations of this pilot study. Firstly, the adult RM model of SHIV rebound used in the study was not originally designed for a direct comparison with the infant ATI model (e.g., infection route, infection dose and weeks of therapy were not precisely matched to the infant study). Yet, the availability of a comparable adult cohort infected with the same SHIV variant and similar viral replication kinetics provided us a unique opportunity to investigate the differences in infant immune responses during and after therapy with respect to adults. We also acknowledge that differences in the challenge route and duration of ART in adult RMs might have limited our ability to directly compare the immune responses in the two age groups. Moreover, relatively small cohort sizes likely contributed to our inability to identify some of the previously established immunologic parameters of viral rebound such as pre-therapy levels of T cell exhaustion markers: Tim-3, Lag-3 and PD-1 (15). Hence, further validation of this model in larger pediatric RM cohorts will be required. Secondly, since our cohorts were subjected to very short duration of ART (8-12 weeks), the measured viral reservoir might not be a reflection of the true persistent reservoir on long-term suppressive ART. Therefore, we have excluded measures of viral reservoir size, which have been identified as predictors of viral rebound in previous adult clinical trials (16–18, 46). Finally, low cell numbers collected, and the comprehensive nature of the study has restricted our ability to measure longitudinal T cell functions, potentially missing T cell function-associated correlates of viral rebound time.

In conclusion, this study validated an oral SHIV-infected pediatric infant RM model of ATI and characterized SHIV replication and humoral immune responses during and post-ATI. Larger and longer-term prospective studies will be needed to further optimize this model and identify a comprehensive set of biomarkers that can reliably predict time to viral rebound. Developing algorithms by combining several surrogate markers of viral rebound could greatly accelerate the process of screening children as candidates for ATI trials, and for development of novel therapeutics for HIV cure research. Additionally, an infant HIV rebound model will be a valuable tool to identify and evaluate the potency of novel therapeutic strategies for attaining functional cure in the context of the maturing infant immune system.

## MATERIALS AND METHODS

### Animal care and study design

Type D retrovirus-, SIV- and STLV-1 free Indian rhesus macaques (RM) (*Macaca mulatta*) were maintained in the colony of California National Primate Research Center (CNPRC, Davis, CA), as previously described (47). Six infant RMs were orally challenged with SHIV.CH505.375H.dCT (22) as described in (26). Briefly, the infant RMs were challenged at 4 weeks of age by bottle feeding 3 times/day for 5 days at a dose of 8.5×10^4^ TCID_50_, to mimic breast milk transmission. After one week of challenges, 1 infant became infected. The remaining 5 were sedated and orally challenged weekly at a dose of 6.8×10^5^ TCID_50_. Within three weeks, 4 more became infected and the final one was challenged at increasing doses (1.3×10^6^ TCID_50_, followed by 3.4×10^6^ TCID_50_) until infected at 14 weeks of age (Table 1). Six adult RMs (age between 4-10 years), were intravenously infected with SHIV.CH505.375H.dCT at a dose of 3.4×10^5^ TCID_50_ as described in (26). The plasma viral RNA load of the monkeys were assessed by highly sensitive qRT-PCR (48). A co-formulation containing 5.1 mg/kg tenofovir disoproxil fumarate (TDF), 40 mg/kg emtricitabine (FTC) and 2.5 mg/kg dolutegravir (DTG) was prepared as described previously (49) and administered once daily by the subcutaneous route starting 8 weeks (infants) or 12 weeks (adults), post-infection (w.p.i).

### Collection and processing of blood and tissue specimens and MHC typing of animals

Animals were sedated with ketamine HCl (Parke-Davis) injected at 10 mg/kg body weight. EDTA-anticoagulated blood was collected via peripheral venipuncture. Plasma was separated from whole blood by centrifugation. Tissues were either fixed in formalin for *in situ* hybridization, or mononuclear cells were isolated from the tissues by density gradient centrifugation as described in (37). DNA extracted from splenocytes was used to screen for the presence of the major histocompatibility complex (MHC) class I alleles Mamu-A*01 and -B*01 and -B*08, using a PCR-based technique (50, 51).

### CD4+ T cell subpopulation sorting and cell-associated SHIV DNA and RNA quantification

CD4^+^ T cells were enriched from PBMCs and tissue mononuclear cells using a negative selection magnetic-activated cell sorting (MACS) system as per manufacturer’s instructions (Miltenyi Biotec, Germany). Enriched CD4^+^ T-cells were stained with fluorescently conjugated antibodies listed in Table S2, and sorted for naïve, memory and Tfh CD4+ T cell subsets (Figure S4). Total RNA and genomic DNA were isolated using the RNeasy Mini Kit and DNeasy Blood and Tissue Kit, respectively (Qiagen, Germany). Viral cDNA was generated from the extracted total RNA using SuperScript III reverse transcriptase enzyme (Invitrogen, Carlsbad, CA), PCR nucleotides (NEB, MA) and Gag-specific reverse primer (Table S3). SHIV-DNA and RNA/million CD4+ T cells in blood and necropsy tissues were estimated by amplifying cDNA and genomic DNA with primers and probes described in Table S3, using digital droplet PCR (ddPCR) as described in (37). The sensitivity of the ddPCR assay was estimated to be detection of 1 SHIV gag copy in 10,000 uninfected CD4+ T cells. Therefore, an input CD4+ T cell count of 10,000 was defined as threshold cell count (TCC) for the analysis, and samples having input cells< TCC were not analyzed. SHIV DNA/million CD4+ T cells was estimated after normalizing SHIV gag copy numbers with input CD4+ counts. Quantification of SHIV DNA in peripheral lymph node-associated naïve, memory and Tfh CD4+ T cell population was performed by qPCR as described in (21), using primers and probes described in Table S3.

### Measurement of HIV Env-specific antibody responses by Enzyme-Linked Immunosorbent Assay (ELISA)

The plasma concentrations of HIV Env-specific antibodies were estimated by ELISA, as previously described (29). Human CH22 monoclonal antibody was used as standard and the concentration of HIV Env-specific IgG antibody was calculated relative to the standard using a 5-parameter fit curve (SoftMax Pro 7). The positivity cut off for the assay was defined as the optical density (OD) of the lowest-concentration rhesus IgG standard that was greater than three times the average OD of blank wells. CD4 blocking ELISAs were done as described in (47).

### HIV-1 Env-specific IgG epitope specificity and breadth using binding antibody multiplex assay (BAMA)

HIV antigens were covalently conjugated to polystyrene beads (Bio-Rad), and binding of IgG to the bead-conjugated HIV-1 antigens was measured in RM plasma samples (29). The antigens used for the assay have been described in (26). Purified IgG from pooled plasma of HIV-1 vaccinated macaques (RIVIG) was used as positive control.

### Neutralization assays

Neutralization of MW965.LucR.T2A.ecto/293T IMC (clade C, tier 1) and the autologous CH505.TF (clade C, tier 2) HIV-1 pseudovirus by plasma antibodies was measured in TZM-bl cells as previously described (29, 52, 53). The ID_50_ was calculated as described in (26). The monoclonal antibody, b12R1, was used as a positive control for MW965 virus, and VRC01 was used a positive control for CH505 T/F virus.

### ADCC-GranToxiLux (GTL) Assay

The ADCC-GTL assay was used to measure plasma ADCC activity as previously described (47, 54). CEM.NKR_CCR5_ target cells were coated with recombinant CH505 gp120. Adult and infant plasma samples were tested after a 4-fold serial dilution starting at 1:100.

### Single Genome Amplification (SGA)

SGA of plasma virus was done as previously described (55). Primer used for cDNA preparation was SHIVEnv.R3out. A first round of PCR amplification was conducted using primers SIVmac.F4out and SHIVEnv.R3out A second round of PCR was conducted using primers SIVmac766.F2in and SIVmac766.R2in (Table S3). *Env* gene amplicons obtained were sequenced by Sanger sequencing, and phylogenetic tree were constructed for the aligned *env* gene sequences by neighbor-joining method using Seaview (56). Average pairwise distance was calculated using the MEGA6 software (57).

### Tissue-associated infectious viral titers by co-culture assay

Serial dilutions of RM lymphoid and gut-associated mononuclear cells were co-cultured with Tzm-bl reporter cells as described previously (37) (Figure S2). Fifty percent cellular infectious dose (CID_50_) was calculated as the number of mononuclear cells per 10^4^ mononuclear cells required to yield detectable infection of 50% Tzm-bl cells, using Reed-Muench method. The detection threshold of the assay was established as 2.5 times the mean luminescence output of Tzm-bl only cells from 10 independent experiments (876 RLU).

### *In situ* hybridization (ISH)

Formalin fixed, paraffin embedded tissue sections were seqentially cut (5 µm) and stained for CD3 and CD20 (Table S2) as previously described (58, 59). SHIV RNA was visualized with the 1-Plex ViewRNA^TM^ ISH Tissue Assay Kit using SIV_mac239_ or Beta actin (positive control) probe sets and ViewRNA Chromogenic Signal Amplification Kit (ThermoFisher, Waltham, MA). These two sequential slides were indivudually imaged with a Zeiss AxioObserver microscope and AxioCam MRm camera. Composite overlays of CD3/CD20 stained slides with ISH slides were prepared using Zen Lite v2.3 software (Zeiss).

### T cell phenotyping

Phenotyping of rhesus PBMCs and tissue-associated mononuclear cells were performed as described previously (37). Fluorescently-conjugated antibodies used to stain cells are reported in Table S2. For intercellular staining, cells were fixed and permeabilized using eBioscience^TM^ FoxP3/transcription factor staining buffer set (Thermo Fisher Scientific), according to manufacturer’s instructions. The stained cells were acquired on an LSRII flow cytometer (BD Biosciences) using BD FACS Diva software and analyzed with FlowJo software version 10 (Tree Star, Inc). Gating for all surface and intracellular markers (Figure S4) was based on fluorescence minus one (FMO) controls. Complete blood counts (CBC) were performed on EDTA-anticoagulated blood samples. Samples were analyzed using a Pentra 60C+ analyzer (ABX Diagnostics). Absolute lymphocyte counts in blood were calculated using the PBMC counts obtained by automated complete blood counts, multiplied by the lymphocyte percentages.

### Multiplex analysis of plasma cytokines and chemokines

Plasma cytokine and chemokine concentrations were assayed in duplicate using a MIP-1α singleplex kit and a 22-analyte multiplex panel (G-CSF, GM-CSF, IFN-γ, IL-1RA, IL-1β, IL-2, IL-4, IL-5, IL-6, IL-8, IL-10, IL-12/23 (p40), IL-13, IL-15, IL-17, IL-18, MCP-1, MIP-1β, sCD40L, TGF-α, TNF-α, VEGF) (both Millipore PRCYTOMAG-40K) performed according to the manufacturer’s recommended protocol and read using a FlexMAP 3D array reader (Luminex Corp.). Data were analyzed using Bio-Plex Manager software (Bio-Rad).

### Statistical analysis

Differences in immune assay measurements between pre-identified time points post-infection were performed by Wilcoxon signed-rank test using the R language and environment for statistical computing (60). Four responses were pre-selected as primary variables of interest. Additionally, 5 virologic and humoral response-associated variables and 16 T cell phenotype and plasma inflammatory response-associated variables were selected for exploratory analysis. Cox proportional hazards modeling was used to create univariate survival models for each variable, treating time to rebound as the event of interest. To account for differences between adult and neonate groups, a binary indicator for the two different groups was used as a control variable. To identify correlates of viral rebound in infants and adults, Kendall’s tau between every variable of interest and the time to virus rebound was computed. Raw p-values and/or FDR-adjusted (Benjamini-Hochberg) p-values, and hazard ratios with confidence intervals were reported for each variable.

## ACKNOWLEDGEMENTS

The work was supported by National Institutes of Health grants P01 AI117915 (S.R.P. and K.D.P.); 5R01 AI106380 (S.R.P.); T32 5108303 (A.D.C.) and 5R01-DE025444 (S.R.P.). This work was also supported by the Penn Center for AIDS Research Viral and Molecular Core P30 AI045008 (K.J.B.); the BEAT-HIV: Delaney Collaboratory to Cure HIV-1 Infection by Combination Immunotherapy UM1AI126620 (K.J.B.); and CARE: Delaney Collaboratory for AIDS Eradication UM1AI126619 (K.J.B.) and the Office of Research Infrastructure Program/OD (P51OD11107; CNPRC). The research contributions by A.D.C. and K.D.P. were supported by the University of North Carolina at Chapel Hill Center for AIDS Research (CFAR), and NIH funded program P30 AI050410. The funders had no role in study design, data collection and interpretation, or the decision to submit the work for publication. The content is solely the responsibility of the authors and does not necessarily represent the official views of the National Institutes of Health.

Several protein antigens for BAMAs and ELISAs were generously provided by Dr. Barton Haynes, supported by NIH NIAID Division of AIDS UM1 grant AI100645 for the Center for HIV/AIDS Vaccine Immunology-Immunogen Discovery (CHAVI-ID), and produced at the Duke Human Vaccine institute (DHVI) Protein Production Facility. We thank Dr. Jeff Lifson, Rebecca Shoemaker and colleagues in the Quantitative Molecular Diagnostics Core of the AIDS and Cancer Virus Program of the Frederick National Laboratory for expert assistance with viral load measurements. We would like to thank Jennifer Watanabe, Dr. Amir Ardeshir and the staff of the CNPRC Colony Research Services for their support in these studies. We thank Emory University Pediatrics/Winship Flow Cytometry Core for technical assistance with CD4+ T cell subpopulation sorting and Center for AIDS Research at Emory University (P30AI050409) for qPCR assays. We thank Papa Kwadwo Morgan-Asiedu, Joshua A. Eudailey, Holly Heimsath and Shuk Hang (Grace) Li at DHVI for assisting with animal tissue processing. We would also like to thank DHVI Regional Biocontainment laboratory (RBL) for technical support with multiplex assay for measuring plasma cytokine response, and R. Whitney Edwards and Nicole Rodgers for performing ADCC assays. T cell phenotyping was performed in the DHVI Flow Cytometry Facility (Durham, NC).

## AUTHOR DISCLOSURE STATEMENT

No competing financial interests.

## AUTHOR CONTRIBUTIONS

R.G., A.N.N., K.D.P., K.K.A.V.R. and S.R.P. designed the study and interpreted the data; R.G., A.N.N., J.J.T., M.D., A.K., J.M., R.J.M., C.M., A.D.C., V.O.P., M.V. and J.P. performed the experiments and analyzed the data; L.F. and C.C. performed the statistical analysis; G.M.S. and K.J.B. provided virus and consulted on the infection model; A.C., K.D.P. and K.K.A.V.R. contributed important insights for the interpretation and discussion of the results. R.G. and S.R.P. drafted the manuscript. All authors read and approved the final manuscript.

## SUPPLEMENTAL MATERIALS

**Figure S1: Proportions of CD4+ T cells in blood, lymph nodes and gut-associated tissues of SHIV.CH505.375H.dCT infected infants and adult RMs. (A)** Frequencies of PBMC-CD4+ T cells in SHIV.CH505.375H.dCT infected infant and adult RMs through 28 w.p.i and 32 w.p.i, respectively. **(B)** Proportions of CD4+ T cells of CD3+ T cells in PBMC, oral and gut-associated lymphoid tissues, and spleen of infected infant RMs at necropsy (28 w.p.i) and adult RMs at necropsy (32 w.p.i). Red symbols represent infants and blue symbols represent adults. Each symbol represents one animal. Yellow and grey boxes represent duration of ART and duration of ATI, respectively. Infants with plasma VL<15 copies/mL at 12 w.p.i has been represented with open symbols.

**Figure S2: Tissue-associated infectious viral levels in RMs infected with SHIV.CH505.375H.dCT.** Mononuclear cells isolated from tissues of **(A)** infant and **(B)** adult RMs infected with SHIV.CH505.375H.dCT were serially diluted and co-cultured with Tzm-bl reporter cells for 72 hrs, followed by luminescent detection of tissue-associated SHIV infectivity in relative luminescence units (RLU). The RLU limit of detection for positive tissue-associated SHIV infection (dashed line) was defined as 2.5 times the mean maximum RLU elicited form Tzm-bl cells (n=10 independent assays) in the co-culture assay.

**Figure S3: Specificity of Env-IgG responses before- and on-ART in SHIV.CH505.375H.dCT infected RMs.** Plasma IgG specificity against a panel of HIV Env linear and conformational epitopes pre ART (12 w.p.i) and on ART (20 w.p.i) in SHIV.C.CH505 infected **(A)** infant and **(B)** adult RMs. Heat maps represents mean fluorescence intensity (MFI) of IgG binding to each epitope.

**Figure S4: Flow cytometry gating strategy for T cell phenotyping and sorting.** For T cell phenotyping, CD4+ T cells and CD8+ T cells were positively selected from the PBMCs by sequential selection of forward and side scatter singlets, lymphocytes, viable cells, CD16-CD14-(Monocytes/macrophages) cells and CD3+ cells (T cells). CD4+ T cells were further analyzed for expression of activation markers HLA-DR and CD69; proliferation marker Ki67 and exhaustion markers PD-1. For sorting CD4+ T cells, CD4+ T cells were positively selected from lymph node-associated mononuclear cells by sequential selection of forward scatter singlets, lymphocytes and CD3^+^ T cells. Tfh cells (CXCR5^hi^ PD-1^hi^), naïve CD4+ T cells (CD95-CD28+ CD45RA+ CCR7+) and memory CD4+ T cells (CD95+ CD28+ CD45RA-CCR7+) were sorted from CD4+ T cells.

**Table S1.** MHC class I genotype of infant and adult RMs.

**Table S2.** Antibodies used for T cell phenotyping, CD4+ T cell sorting and *in situ* hybridization (ISH).

**Table S3.** Primers and probes used for the assays.

