## Supplemental materials for "Analytical treatment interruption after short-term anti-retroviral therapy in a postnatally SHIV infected infant rhesus macaque model"

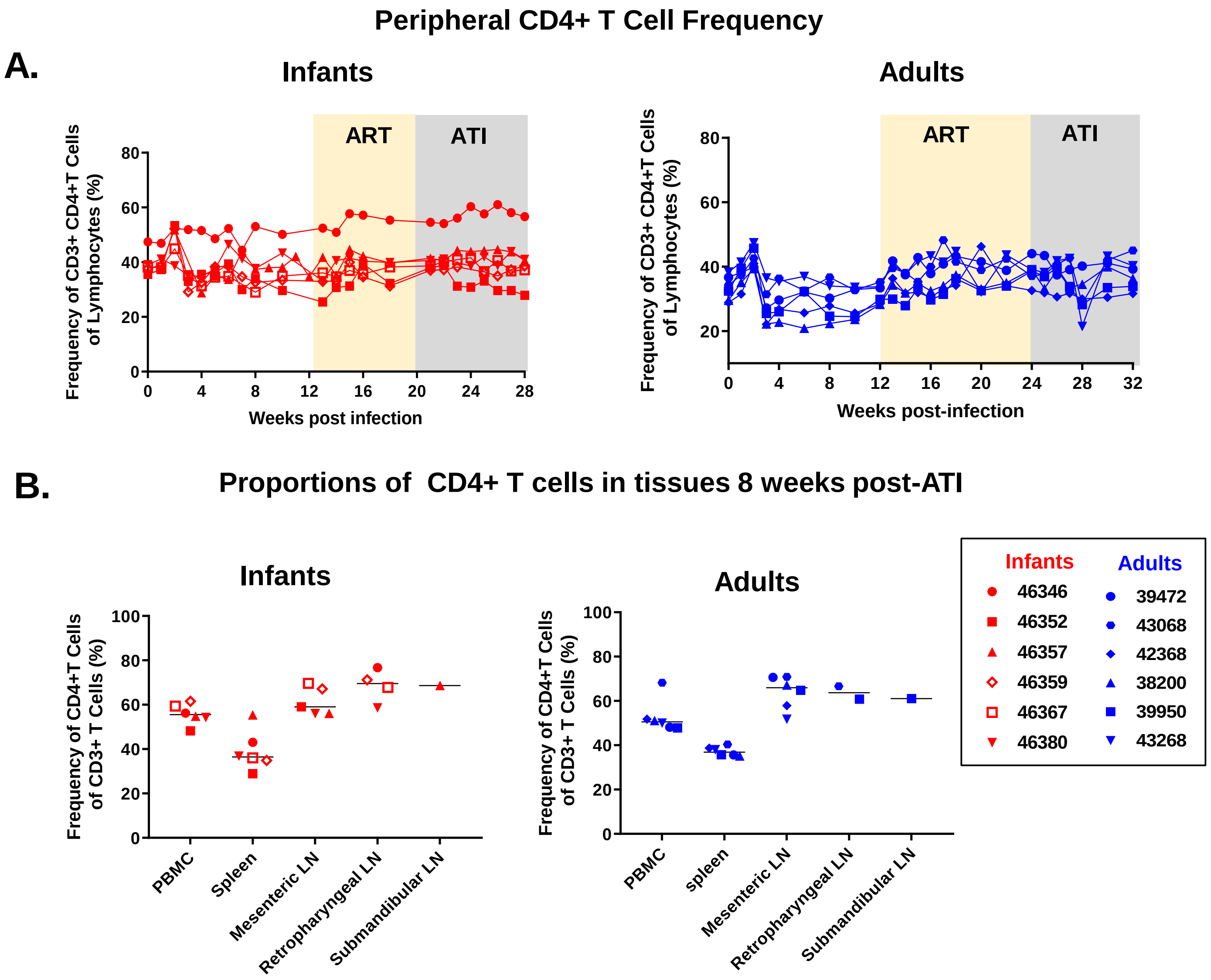


**Figure S1: Proportions of CD4+ T cells in blood, lymph nodes and gut-associated tissues of SHIV.CH505.375H.dCT infected infants and adult RMs. (A)** Frequencies of PBMC-CD4+ T cells in SHIV.CH505.375H.dCT infected infant and adult RMs through 28 w.p.i and 32 w.p.i, respectively. **(B)** Proportions of CD4+ T cells of CD3+ T cells in PBMC, oral and gut-associated lymphoid tissues, and spleen of infected infant RMs at necropsy (28 w.p.i) and adult RMs at necropsy (32 w.p.i). Red symbols represent infants and blue symbols represent adults. Each symbol represents one animal. Yellow and grey boxes represent duration of ART and duration of ATI, respectively. Infants with plasma VL<15 copies/mL at 12 w.p.i has been represented with open symbols.

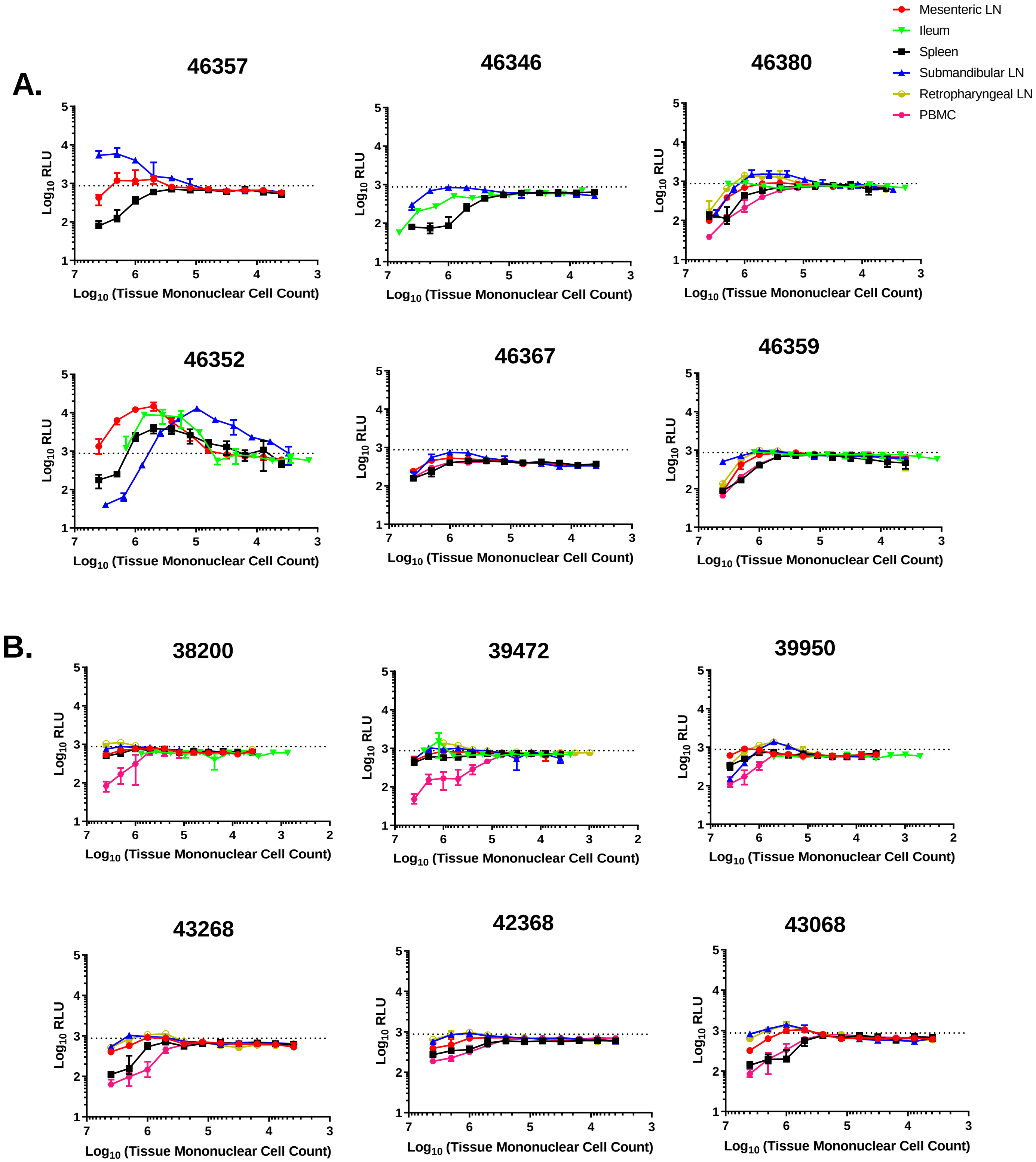

A.
46357
46346
46380
46352
46367
46359
B.
38200
39472
39950
43268
42368
43068
$Log_{10}$ RLU
$Log_{10}$ (Tissue Mononuclear Cell Count)
Mesenteric LN
Ileum
Spleen
Submandibular LN
Retropharyngeal LN
PBMC

**Figure S2: Tissue-associated infectious viral levels in RMs infected with SHIV.CH505.375H.dCT.** Mononuclear cells isolated from tissues of **(A)** infant and **(B)** adult RMs infected with SHIV.CH505.375H.dCT were serially diluted and co-cultured with Tzm-bl reporter cells for 72 hrs, followed by luminescent detection of tissue-associated SHIV infectivity in relative luminescence units (RLU). The RLU limit of detection for positive tissue-associated SHIV infection (dashed line) was defined as 2.5 times the mean maximum RLU elicited form Tzm-bl cells (n=10 independent assays) in the co-culture assay.

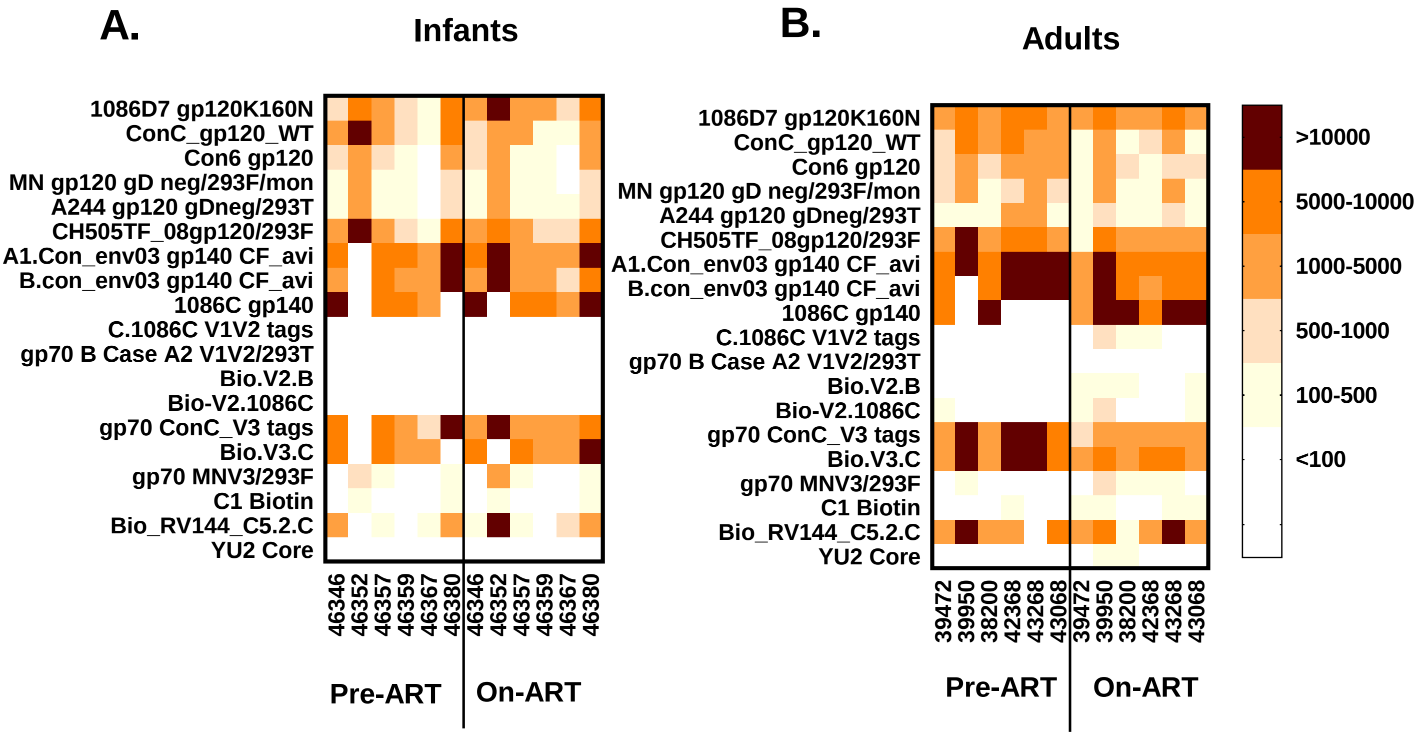


**Figure S3: Specificity of Env-IgG responses before- and on-ART in SHIV.CH505.375H.dCT infected RMs.** Plasma IgG specificity against a panel of HIV Env linear and conformational epitopes pre ART (12 w.p.i) and on ART (20 w.p.i) in SHIV.C.CH505 infected **(A)** infant and **(B)** adult RMs. Heat maps represents mean fluorescence intensity (MFI) of IgG binding to each epitope.

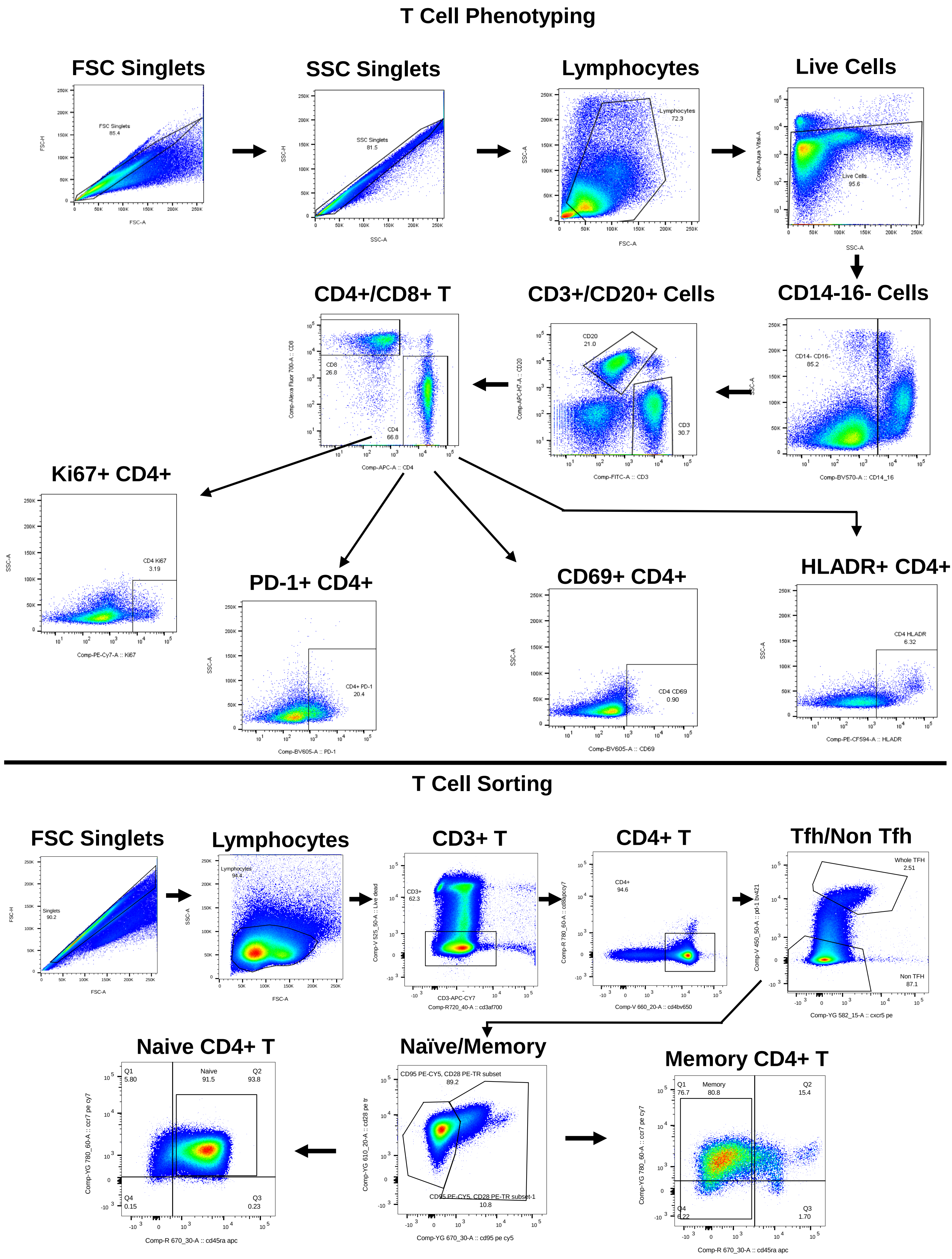

T Cell Phenotyping
FSC Singlets
FSC Singlets 85.4
SSC Singlets
SSC Singlets 81.5
Lymphocytes
Lymphocytes 72.3
Live Cells
Live Cells 95.6
CD14-16- Cells
CD14- CD16- 85.2
CD3+/CD20+ Cells
CD20 21.0
CD3 30.7
CD4+/CD8+ T
CD8 26.8
CD4 66.8
Ki67+ CD4+
CD4 Ki67 3.19
PD-1+ CD4+
CD4+ PD-1 20.4
CD69+ CD4+
CD4 CD69 0.90
HLADR+ CD4+
CD4 HLADR 6.32
T Cell Sorting
FSC Singlets
Singlets 90.2
Lymphocytes
Lymphocytes 94.4
CD3+ T
CD3+ 62.3
CD4+ T
CD4+ 94.6
Tfh/Non Tfh
Whole TFH 2.51
Non TFH 87.1
Naïve/Memory
CD95 PE-CY5, CD28 PE-TR subset 89.2
CD95 PE-CY5, CD28 PE-TR subset-1 10.8
Naive CD4+ T
Q1 5.80
Naive 91.5
Q2 93.8
Q4 0.15
Q3 0.23
Memory CD4+ T
Q1 76.7
Memory 80.8
Q2 15.4
Q4 6.22
Q3 1.70

**Figure S4: Flow cytometry gating strategy for T cell phenotyping and sorting.** For T cell phenotyping, CD4+ T cells and CD8+ T cells were positively selected from the PBMCs by sequential selection of forward and side scatter singlets, lymphocytes, viable cells, CD16- CD14- (Monocytes/macrophages) cells and CD3+ cells (T cells). CD4+ T cells were further analyzed for expression of activation markers HLA-DR and CD69; proliferation marker Ki67 and exhaustion markers PD-1. For sorting CD4+ T cells, CD4+ T cells were positively selected from lymph node-associated mononuclear cells by sequential selection of forward scatter singlets, lymphocytes and $CD3^{+}$ T cells. Tfh cells ($CXCR5^{hi}$ $PD\text{-}1^{hi}$), naïve CD4+ T cells (CD95- CD28+ CD45RA+ CCR7+) and memory CD4+ T cells (CD95+ CD28+ CD45RA- CCR7+) were sorted from CD4+ T cells.

**Table S1.** MHC class I genotype of infant and adult RMs.

| | Animal ID | MHC Class I[a] | | |
|---|---|---|---|---|
| | | Mamu A | | Mamu B |
| | | A*01 | A*08 | B*01 |
| Infant | 46357 | - | - | +/- |
| | 46346 | - | - | - |
| | 46352 | - | +/- | +/- |
| | 46359 | - | - | - |
| | 46367 | - | - | - |
| | 46380 | - | - | - |
| Adult | 39472 | - | - | |
| | 43068 | - | - | +/- |
| | 43268 | - | - | +/- |
| | 42368 | - | - | - |
| | 39950 | - | - | - |
| | 38200 | - | - | +/+ |

[a]Mamu A*01 has been associated with attenuation of SHIV disease progression and Mamu B*01 has been associated with better SIV disease progression.

+/- indicate heterozygous.

**Table S2.** Antibodies used for T cell phenotyping, CD4+ T cell sorting and *in situ* hybridization (ISH).

| | Marker | Fluorophore | Staining type | Clone | Vendor | Secondary Antibody |
|---|---|---|---|---|---|---|
| T Cell Phenotyping | CD20 | APC-H7 | Surface | 2H7 | BD Biosciences | N/A |
| | CD3 | FITC | Surface | SP34 | BD Biosciences | N/A |
| | CD4 | APC | Surface | L200 | BD Biosciences | N/A |
| | CD8 | AF700 | Surface | SK1 | BD Biosciences | N/A |
| | Ki67 | PEcy7 | Intracellular | B56 | BD Biosciences | N/A |
| | HLADR | PE-CF594 | Surface | G46-6 | BD Biosciences | N/A |
| | CD69 | BV605 | Surface | FN50 | BD Biosciences | N/A |
| | CD14 | BV570 | Surface | M5E2 | Biolegend | N/A |
| | CD16 | BV570 | Surface | 3G8 | Biolegend | N/A |
| | PD-1 | BV605 | Surface | EH12.2H7 | Biolegend | N/A |
| CD4+ T Cell Sorting | CD3 | AF700 | Surface | SP34-2 | BD Biosciences | N/A |
| | CCR7 | PE-CY7 | Surface | CD197 | BD Biosciences | N/A |
| | CD8 | APC-CY7 | Surface | SK1 | BD Biosciences | N/A |
| | CD45RA | APC | Surface | 5H9 | BD Biosciences | N/A |
| | CD95 | PE-CY5 | Surface | DX2 | BD Biosciences | N/A |
| | CD28 | ECD | Surface | CD28.2 | Beckman Coulter | N/A |
| | CD4 | BV650 | Surface | OKT4 | Biolegend | N/A |
| | PD-1 | BV421 | Surface | EH12.2H7 | Biolegend | N/A |
| | CXCR5 | PE | Surface | MU5UBEE | eBiosciences | N/A |
| ISH | CD3 | None | Surface | Polyclonal rabbit IgG | Dako | Goat anti-rabbit AF488 (Invitrogen) |
| | CD20 | None | Surface | Mouse IgG2a-L26 | Dako | Goat anti-mouse AF594 (Invitrogen) |

[a]N/A, Not applicable

**Table S3.** Primers and probes used for the assays.

| Primers and Probes | Identifier | Type | Assay |
|---|---|---|---|
| 5’-CTACTGGTCTCCTCCAAAGAGAGAATTG-3’ | Gag-specific reverse primer | Primer | cDNA production |
| 5’-TCAGGCACTGTCAGAAGGTT-3’ | Gag-specific forward primer | Primer | ddPCR |
| 5’ TTGTTGTGGAGCTGGTTGTG-3’ | Gag-specific reverse primer | Primer | ddPCR |
| 5’-FAM-AGCCGCTTGATGGTCTCCCACA-TAMRA-3’ | SHIV CH505 gag-specific probe | Probe | ddPCR |
| 5’ -GAAGGTGAAGGTCGGAGTC-3’ | Gag-specific forward primer | Primer | qPCR |
| 5’-GAAGATGGTGATGGGATTTC-3’ | Gag-specific reverse primer | Primer | qPCR |
| 5’ -TGCATGAGAAAACGCCAGTAA-3’ | Albumin-specific forward primer | Primer | qPCR |
| 5’-ATGGTCGCCTGTTCACCAA-3’ | Albumin-specific reverse primer | Primer | qPCR |
| 5’-FAM CAAGCTTCCCGTTCTCAGCC TAMRA-3’ | SIVmac gag-specific probe | Probe | qPCR |
| 5’-6-FAM AGAAAGTCACCAAATGCTGCACGGAATC-3’-6-TAMRASp | Albumin probe | Probe | qPCR |
| 5’-CTAATTCCTGGTCCTGAGGTGTAATCCTG-3’ | SHIVEnv.R3out | Primer | SGA |
| 5’-TCATATCTATAATAGACATGGAGACACCC-3’ | SIVmac.F4out | Primer | SGA |
| 5’- CTAATTCCTGGTCCTGAGGTGTAATCCTG-3’ | SHIVEnv.R3out | Primer | SGA |
| 5’- GGAAATCCTCTCTCAACTATACCGCCCTC -3’ | SIVmac766.F2in | Primer | SGA |
| 5’- CTATTGCCAATTTGTAACTCATTGTTC-3’ | SIVmac766.R2in | Primer | SGA |
